# TGF-β induces matrisome pathological alterations and EMT in patient-derived prostate cancer tumoroids

**DOI:** 10.1101/2023.04.03.534859

**Authors:** Soraia Fernandes, Jorge Oliver-De La Cruz, Marco Cassani, Sofia Morazzo, Helena Ďuríková, Alessio Caravella, Piergiuseppe Fiore, Giulia Azzato, Giuseppe De Marco, Agostino Lauria, Valerio Izzi, Veronika Bosáková, Jan Fric, Petr Filipensky, Giancarlo Forte

**Affiliations:** Center for Translational Medicine (CTM), International Clinical Research Centre (ICRC), St. Annés Hospital, Brno, Czech Republic; Faculty of Medicine, Department of Biomedical Sciences, Masaryk University, 62500, Brno, Czech Republic; Department of Computer Engineering, Modelling, Electronics and Systems Engineering (DIMES), University of Calabria (UNICAL), Via P. Bucci, Cubo 42C, Rende (CS), 87036, Italy; Information Technology Center (ICT), University of Calabria (UNICAL), Via P. Bucci, Cubo 22B, Rende (CS), 87036, Italy; Department of Engineering for Innovation, University of Salento (UNISALENTO),Corpo Z, Campus Ecotekne, SP.6 per Monteroni, Lecce (LE), Italy; Faculty of Biochemistry and Molecular Medicine, University of Oulu, Oulu FI-90014, Finland; Faculty of Medicine, BioIM Research Unit, University of Oulu, Oulu FI-90014, Finland; Foundation for the Finnish Cancer Institute, Tukholmankatu 8, Helsinki, Finland; Institute of Hematology and Blood Transfusion, Prague, Czech Republic; Department of Urology, St. Annés Hospital, Brno, Czech Republic; School of Cardiovascular and Metabolic Medicine & Sciences, King’s College London, London SE5 9NU, UK

**Author notes:** Institute for Bioengineering of Catalonia (IBEC), The Barcelona Institute for Science and Technology (BIST), Barcelona, Spain. TECHFEM S.p.A. – Human and Sustainable Engineering,Via Toniolo, 1/D, Fano (PU), 61032, Italy. Correspondence Giancarlo Forte, School of Cardiovascular and Metabolic Medicine and Sciences (SCMMS), Faculty of Life Sciences and Medicine King’s College London, SE5 9NU, United Kingdom,; Soraia Fernandes, Center for Translational Medicine (CTM) International Clinical Research Center (ICRC) St. Anne’s University Hospital, Studentska 6, Brno, Czech Republic, 62500.

**Keywords:** Cancer, tumour, extracellular matrix (ECM), tumoroids, prostate, epithelial-to-mesenchymal transition (EMT), transforming growth factor (TGF)-β.

## Abstract

Extracellular matrix (ECM) tumorigenic alterations resulting in high matrix deposition and stiffening are hallmarks of adenocarcinomas and are collectively defined as *desmoplasia*. Here, we thoroughly analysed primary prostate cancer tissues obtained from numerous patients undergoing radical prostatectomy to highlight reproducible structural changes in the ECM leading to the loss of the glandular architecture. Starting from patient cells, we established prostate cancer tumoroids (PCTs) and demonstrated they require TGF-β signalling pathway activity to preserve phenotypical and structural similarities with the tissue of origin. By modulating TGF-β signalling pathway in PCTs, we unveiled its role in ECM accumulation and remodelling in prostate cancer. We also found that TGF-β-induced ECM remodelling is responsible for the initiation of prostate cell epithelial-to-mesenchymal transition (EMT) and the acquisition of a migratory, invasive phenotype. Our findings highlight the cooperative role of TGF-β signalling and ECM *desmoplasia* in prompting prostate cell EMT and promoting tumour progression and dissemination

## Introduction

During carcinogenesis, cells undergo epithelial-to-mesenchymal transition (EMT), a complex dynamic process that causes changes in cellular organization by repressing epithelial features while enhancing the acquisition of mesenchymal phenotype.^1^ In solid tumours, the latter is associated with increased cell migration and invasion. EMT is triggered by changes in gene expression and post-translational regulation mechanisms driven in tumour cells by cues generated by the different stromal cell types and the extracellular matrix (ECM) composing the surrounding tumour microenvironment (TME).^1, 2^ The existence of a positive loop by which changes in tumour cells affect TME structure and composition has also been described.^3^

An important hallmark of tumour development and progression is *desmoplasia*, a phenomenon through which ECM acquires fibrotic features.^4, 5^ This event has been well described for some cancer types, *e.g.* breast and pancreatic ductal adenocarcinomas. In breast cancer, increased ECM stiffness correlates with EMT, invasion and metastasis and poor prognosis^6–8^. The occurrence of *desmoplasia* in other cancer types, like prostate carcinoma is still debated, with little data supporting this hypothesis being provided by studies on commercially available prostate cancer cell lines.^9–13, 14, 15^ The critical mechanisms of ECM remodelling in cancer progression include: (1) increased matrix deposition; (2) chemical modification of ECM components; (3) proteolytic degradation (metalloproteinases - MMPs); and (4) structural changes leading to fibres alignment and increased stiffness.^5, 16^ These processes are altogether driven by the activation of both cancer and tumour-associated stromal cells, such as cancer-associated fibroblasts (CAFs).^5^ CAFs are described as the main producers of ECM in solid tumours. They are known to secrete a variety of signals, comprising the transforming growth factor (TGF-β).^4^ Due to its interference with the expression of different genes and proteins related to cytoskeleton assembly, cell-cell attachment and ECM remodelling in all cell types composing the tumour stroma^2, 17^, TGF-β signalling is considered the major determinant of EMT, invasion and metastasis in solid tumours.^16, 18^ Nevertheless, whether carcinoma cells respond to TGF-β by inducing changes in TME is poorly understood.^17^

Given its intrinsic complexity, TME is not easily replicated in 2D *in vitro* models.^19^ Therefore, novel cellular models have been recently developed that might aid our understanding of the mechanisms driving cancer progression. Recently, 3D *in vitro* tumour models, also known as tumoroids or cancer organoids, have been defined as organ miniatures or avatars resembling tumour complexity.^20^ They have gained momentum as fundamental tools to understand the biology of tumour progression and as patient-specific platforms to test drug sensitivity.^21, 22^ Tumoroids can be obtained *ex-vivo* from pathological specimens, thus offering the possibility to develop *in vitro* microtissues featuring the patient-specific genetic background and allowing for the assessment of their responses to drugs, eventually leading to personalized approaches.^23^ While great efforts have been produced to characterize the cellular composition of tumoroids, the role of the ECM in these advanced cellular models has been, so far, poorly described.

Here, we established patient-derived prostate cancer tumoroids (PCTs) that mimic key features of the native TME, namely the pathological signature of tumorigenic ECM remodelling and the capacity to undergo EMT. We proved that PCTs represent patient-specific and histologically relevant 3D cell models endowed with multicellularity, self-organization, and reflect the complexity of the tissue of origin. By using these models, we gained insight into the process of ECM *desmoplasia* in prostate cancer and resolved the structural and mechanical changes induced by TGF-β stimulation and its effects on tumour metastatic dissemination.

## Results

### Increased ECM production and EMT are pathological hallmarks of primary prostate cancer

To investigate the pathological features occurring in prostate cancer (PCa), specimens from tumour tissues and adjacent non-tumour regions were obtained from radical prostatectomy and analysed by histology and immunohistochemistry (IHC). Supplementary Table 1 and Figure S1 summarize the clinical pathophysiological features of the 80 patients from whom the samples used in the study were obtained, including Gleason score, Prostate Specific Antigen (PSA) levels in the blood and disease incidence based on age and body mass index (BMI). Histological examination of the harvested tissues was conducted to confirm the presence of tumorigenic features in the tissue samples, as shown in Figure S2 by haematoxylin and eosin staining (H&E). Distinct pathological signs in the glandular pattern were visible in tumour samples, confirming the disruption of tissue architecture and stromal invasion during disease progression. Furthermore, the differential expression of N-cadherin (N-CAD) and vimentin (VIM), regarded as characteristic mesenchymal signature and established EMT markers, were found highly expressed in tumorigenic regions, while E-cadherin (E-CAD), a protein used to identify epithelial cells, was found expressed in both tumour and adjacent regions (Figure 1A). In addition, reduced expression of integrin alpha-6 (ITGA6) was detected in tumour sections, suggesting the loss of basal membrane, while luminal cells positive for cytokeratin 8 (CK8) could be detected in both tumour and adjacent regions (Figure 1A). Next, we used Masson’s trichrome staining and the colorimetric measurement of hydroxyproline content to see the distribution and quantify collagen in the patients’ tissues. These analyses revealed that tumours had increased collagen content as compared to their adjacent healthy tissue. (Figure 1B, C and Figure S3). These results, which indicate that ECM accumulates at the tumour site, were further confirmed by IHC analysis performed on either the native tissues or after their decellularization (dECMs).^24^ A significant accumulation of fibronectin (FN) could also be confirmed using the same approach (Figure 1D, E).

**Figure 1.**
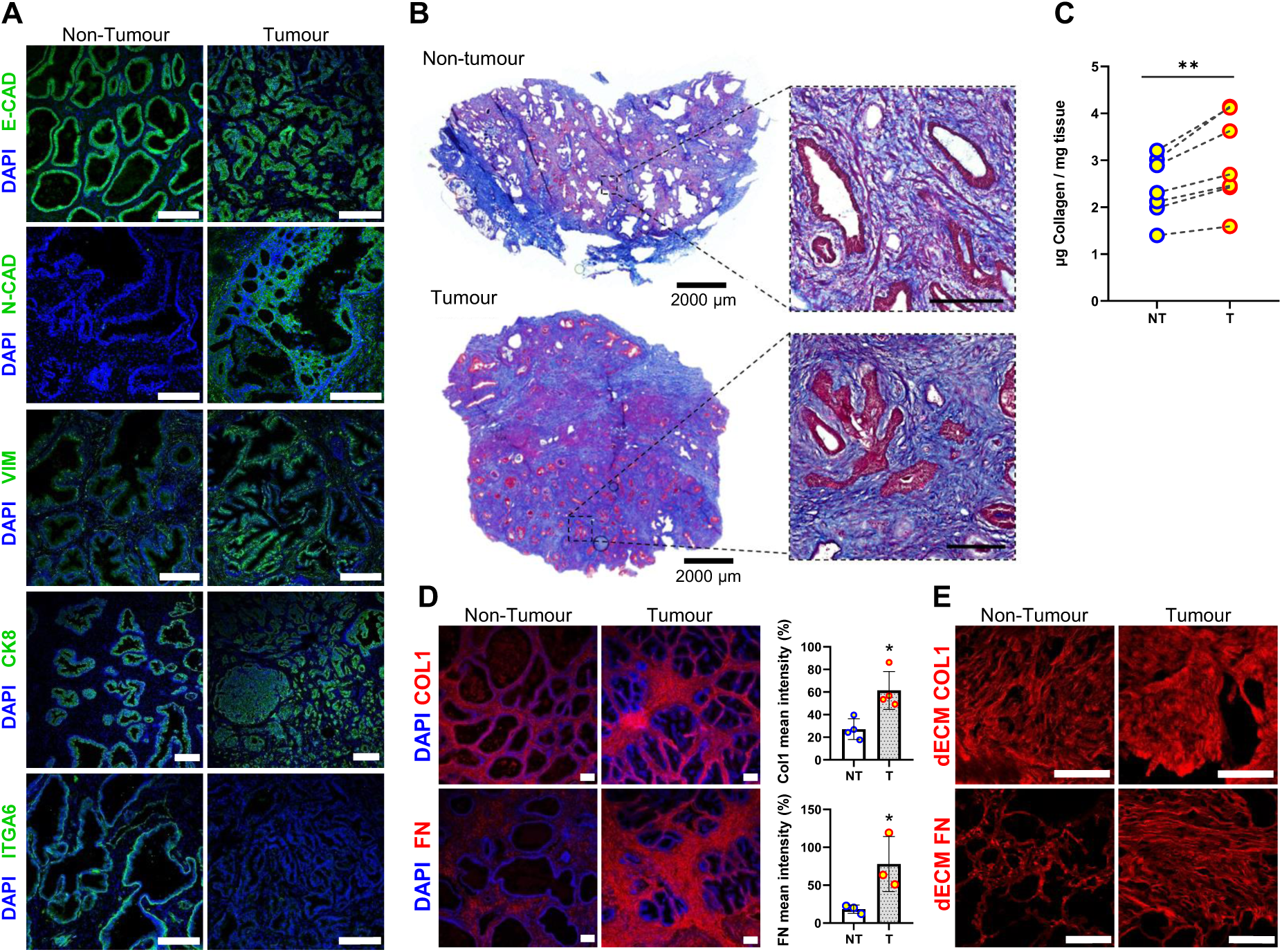
EMT and increased local ECM deposition are distinctive pathological signs of prostate cancer (PCa). **(A)** Representative confocal microscopy images of EMT (E-CAD, N-CAD, VIM), epithelial basal (ITGA6) and luminal markers (CK8) expression in tumour and tumour-adjacent (non-tumour) prostate tissues. Scale bars: 200 µm. **(B)** Representative Masson’s trichrome staining images of tumour and tumour-adjacent prostate tissues. Scale bars: 200 µm. **(C)** Total collagen quantification in tumour and non-tumour adjacent tissue specimens (N = 7); Statistical analysis performed by Wilcoxon matched pairs signed rank test. **(D)** Representative confocal microscopy images of the indicated ECM markers and their relative intensity quantification in tumour and tumour-adjacent prostate tissues. Scale bars: 200 µm. Statistical analysis was performed by Welch’s t-test (n ≥ 3). **(E)** Representative confocal microscopy images of tumour and non-tumour decellularized matrices (dECMs) stained for COL1 and FN. Scale bars: 200 µm. NT: non-tumour; T: tumour.

To gain an insight into ECM deposition in prostate cancer development, we decellularized tumour and tumour adjacent samples and analysed their ultrastructure surface topography using scanning electron microscopy (SEM). The results obtained showed that non-tumorigenic ECM displayed a characteristic structure defined by wavy fibre bundles. On the contrary, tumour specimens exhibited signs of remodelling which led to the formation of a flat, mostly aligned surface. This phenomenon could be quantified by image analysis by parameters like fibre coherency and distribution of orientation (Figure 2A-C and Figure S4).

**Figure 2.**
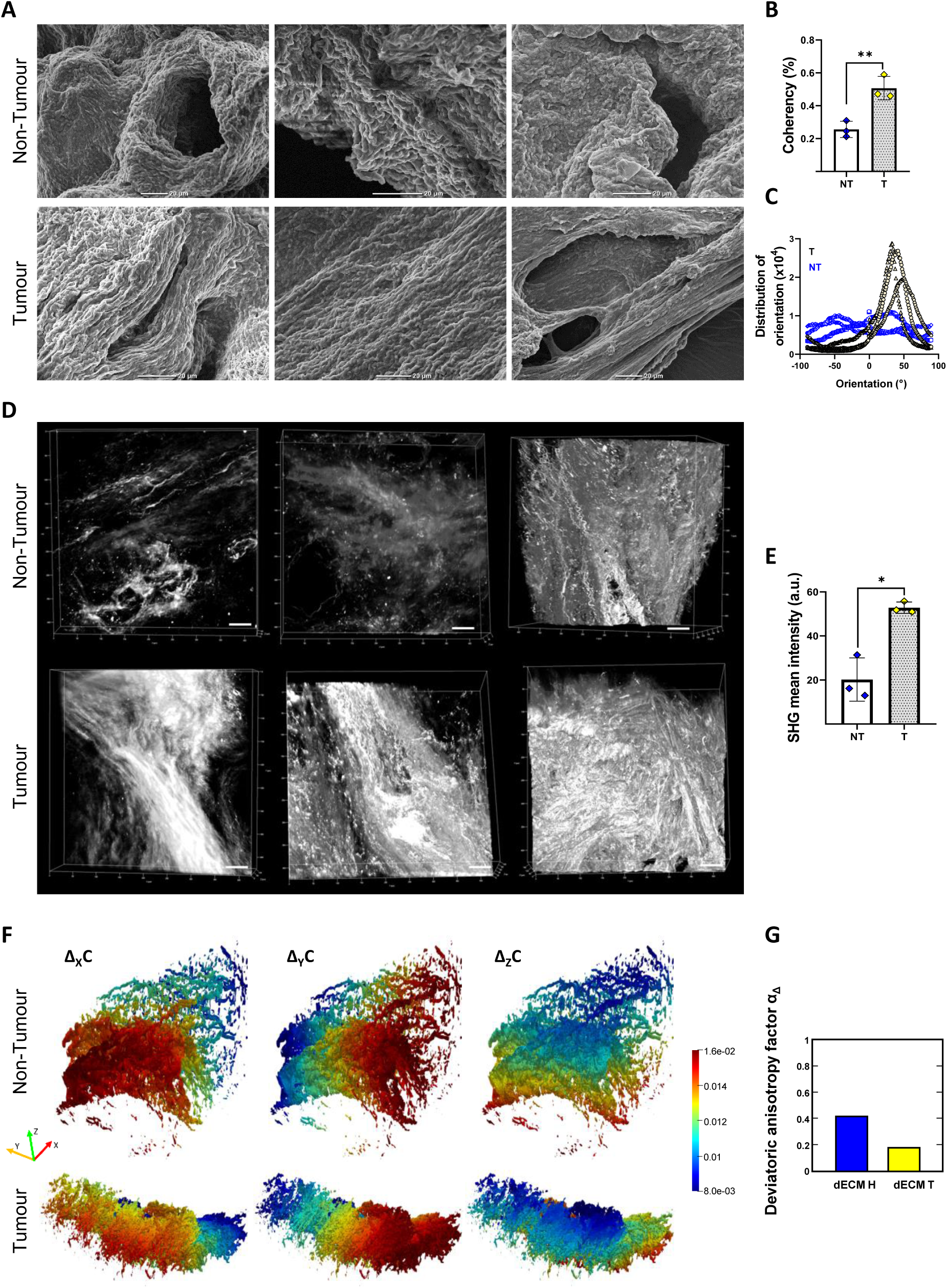
Patient-derived decellularized extracellular matrices (dECMs) reveal consistent structural changes in human prostate cancer. **(A)** Representative scanning electron microscopy (SEM) images of the surface of dECMs obtained from tumour (T) and non-tumour tissues (NT). Scale bars: 20 µm. **(B)** Coherency statistical analysis performed by t-test, N = 3. **(C)** Analysis of fibres orientation in dECMs obtained from 3 different patient samples. **(D-E)** Two-photon second harmonic generation (SHG) representative 3D images of tumour (T) and non-tumour (NT) dECMs and relative SHG mean intensity quantification (N = 3). Scale bars: 50 µm. **(F)** Representative color-coded images of concentration profiles of diffusive surface tortuosity obtained by 3-dimensional (3D) computational fluid dynamics (CFD) simulation of tumour (T) and non-tumour (NT) dECMs. (G) Deviatoric anisotropy (α_Δ_) quantification for tumour and non-tumour dECMs.

Next, we used 2-photon microscopy to resolve the 3D configuration of collagen fibres within the dECMs through second harmonic generation (SHG). Upon 3D reconstruction, tumorigenic dECMs displayed a dense aligned collagen mesh that contrasted with less compact non-tumorigenic dECM (Figure 2D). The compact arrangement of collagen fibres in tumour dECMs determined a 2-fold increment in SHG signal (Figure 2E). Furthermore, starting from their SHG images, two samples derived from the same patient from a tumour and an adjacent non tumour region were characterized by an innovative computational fluid dynamics (CFD) approach allowing for the evaluation of anisotropy from their digitally reconstructed structures. Using this methodology, we assessed the structure anisotropy by evaluating the average effective diffusivity tensor, to obtain tortuosity and connectivity tensors and ultimately the deviatoric anisotropy factor α_Δ_.^25^ This factor subtracts the minimum tortuosity from the overall tortuosity tensor, resulting in a more sensitive estimation of the structural anisotropy.^25^ Representative color-coded images of the concentration profiles calculated by CFD (Figure 2F) and the estimated deviatoric anisotropy factor α_Δ_ (Figure 2G) showed that non-tumour adjacent tissues displayed a more constant topology, *i.e.* similar properties in all positions, while tumour tissues appeared to be less uniform. The results from the CFD also showed a slight decrease in tissue porosity for the tumour-affected ECM (tumour: 0.9844 *vs* non-tumour: 0.9053) which most likely is related to the appearance of desmoplastic features with loss of glandular differentiation. Altogether, these results revealed significant alterations in the ECM organization of primary prostate tissue which occurs alongside EMT and the loss of basal layer integrity.

### Generation and characterization of patient-derived prostate cancer tumoroids (PCTs) from primary PCa tissue

Next, we generated patient-derived prostate cancer tumoroids (PCTs) to investigate the molecular mechanisms leading to ECM accumulation and remodelling and its relationship with the occurrence of EMT in PCa. PCa tissues collected after radical prostatectomy were prepared according to a procedure reported in the literature, with minor modifications (Figure 3A).^26–28^ Briefly, the tissues were first minced into small pieces and then enzymatically digested in a rotatory tube overnight at 37 °C. Next, single cell suspension was seeded in ultra-low attachment plates using supplemented medium cocktail DMEM/F12++ (Table 2). As soon as 48 hours after seeding, floating tumoroids could be visualized (Figure 3A). The procedure reproducibility allowed us to generate tumoroids from all 80 patients enrolled in the study independently of their pathological features and tumour grading (Figure 3B; Supplementary Table 1). We thus interrogated the histological characteristics and cell morphology of the PCTs by H&E and IHC and validated them against their native tissues. The tumoroids displayed variable shapes as visible from H&E and bright field microscopy images (Figure 3C), with irregular acinar morphology originated from poorly differentiated glands. We stained the PCTs with specific antibodies directed against epithelial (E-CAD) and mesenchymal proteins (VIM) and found that both cell types representative of the glandular tissue composition could be found in the tumoroids (Figure 2D). Vimentin was mostly expressed at the edges of the tumoroids, while the core was composed mainly of E-cadherin-positive epithelial cells. Consistent with their histological resemblance to PCa tissues, the epithelial component of the tumoroids was mainly represented by CK8-positive luminal cells (Figure 3E, 5 days of culture) which are the most represented component in cancer tissue (see Figure 1A). Overall, the PCTs recapitulated the given features of the original tumour tissue, including histological and structural resemblance (Figure 3C) and multicellular configuration (Figure 3D).

**Figure 3.**
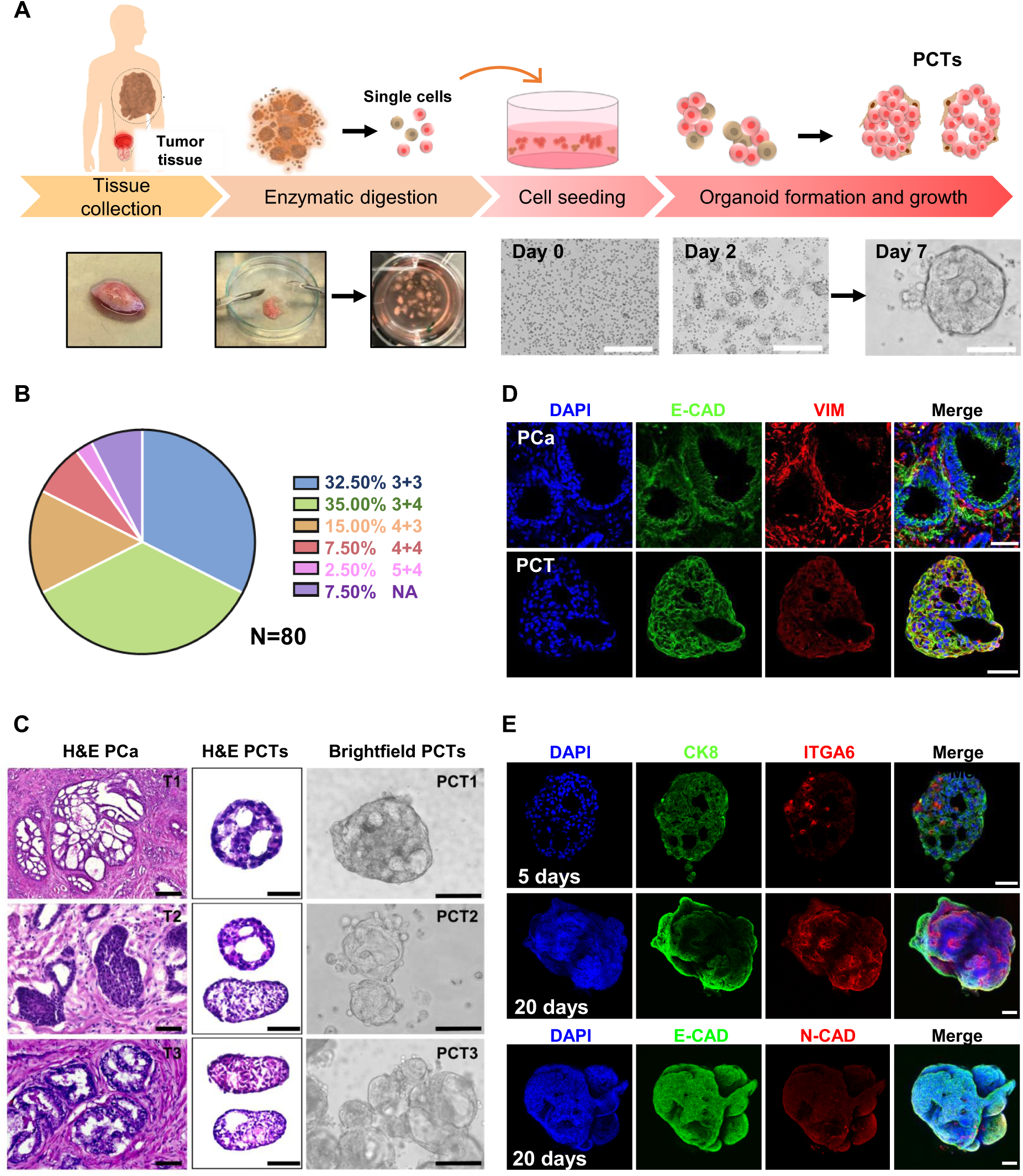
Generation of prostate cancer tumoroids (PCTs) from patient tissues. **(A)** Schematic representation of the procedure used to develop the 3D tumoroids after tissue collection from radical prostatectomy. Scale bars: 100 µm. **(B)** Pie chart representation of the percentage of samples used to generate the PCTs (N = 80 patients) clustered by Gleason Score (GS). **(C)** Representative H&E (left) and bright field (right) images of native tumour tissues and the tumoroids derived from them. Scale bars: 100 µm. **(D)** Representative confocal microscopy images of the PCTs stained for epithelial (E-CAD) and mesenchymal (VIM) markers. Scale bars: 50 µm. **(E)** Representative confocal microscopy images of the PCTs stained for the specified markers at the indicated time in culture. Scale bars: 50 µm.

However, as recently described by Servant *et al,*^28^ over time in culture the prostate tumoroids acquired increased expression of ITGA6-positive cells typical of the basal layer, with benign-like cells overgrowing the tumour ones (Figure 3E). Additionally, low expression of the distinctive N-cadherin (N-CAD) EMT marker was detected after 20 days in culture (Figure 3E). Also, mass spectrometry (MS) analysis confirmed no significant difference could be found in the protein composition of 7 days culture tumoroids from tumour and non-tumour adjacent tissues (N = 4, Figure S5). These data indicated that the culture conditions used favoured the growth of benign-like epithelial cells over their tumour counterpart. These observations underlined the absence of tumorigenic features of the PCTs, typically found in the PCa tissues (Figure 1A) prompting us to explore the possibility of preserving their malignant phenotype.

### TGF-β induces transcriptional changes in PCTs that recapitulate the ECM tumorigenic alterations in PCa

Given the increasing evidence on TGF-β role as tumour promoter,^29^ we hypothesized that a component of the culture medium and a well-known inhibitor of SMAD signalling and EMT induced by TGF-β activity – A83-01 – ^30^could be responsible for the loss of the pathological features in cultured PCTs. To confirm our hypothesis, we conducted a pilot experiment in which we stimulated healthy human prostate cell line (PNT2) with TGF-β (10 µg/mL) for 2 days and analysed the changes in their phenotype. As expected, exposure to TGF-β induced enhanced nuclear expression of its effector SMAD2/3 in PNT2 cells, but also determined the *de novo* expression of mesenchymal markers vimentin and alpha smooth muscle actin (α−SMA, Figure S6). This result was confirmed by flow cytometry, which showed the enhanced expression of vimentin and the concomitant reduction in epithelial marker E-cadherin in cells treated with TGF-β (Figure S7). These findings suggest that the activation of TGF-β canonical pathway, through SMAD2/3 translocation to the nucleus, might trigger EMT in prostate cells, resulting in the amplified expression of mesenchymal markers.

We next moved to investigate the effects of TGF-β pathway activation on PCTs by culturing them with or without TGF-β receptor inhibitor (A83-01). The depletion of the factor from the culture resulted in a significant increase in TGF-β release by the PCTs, as shown by ELISA (Figure S8A). On the contrary, we could not detect any significant change in PSA secretion, which ensured the identity of prostate tissues was not affected by TGF-β stimulation (Figure S8B). We could also confirm that the growth factor activity resulted in an upregulation of SMAD2/3 phosphorylation (Figure 4A) and was associated with the expression of mesenchymal markers, like vimentin (Figure 4B and S9) and α−SMA (Figure 4C) and changed the ratio between N- and E-cadherin in favour of the former (Figure 4D). Together with VIM, α−SMA and N-CAD are established EMT markers.^3^

**Figure 4.**
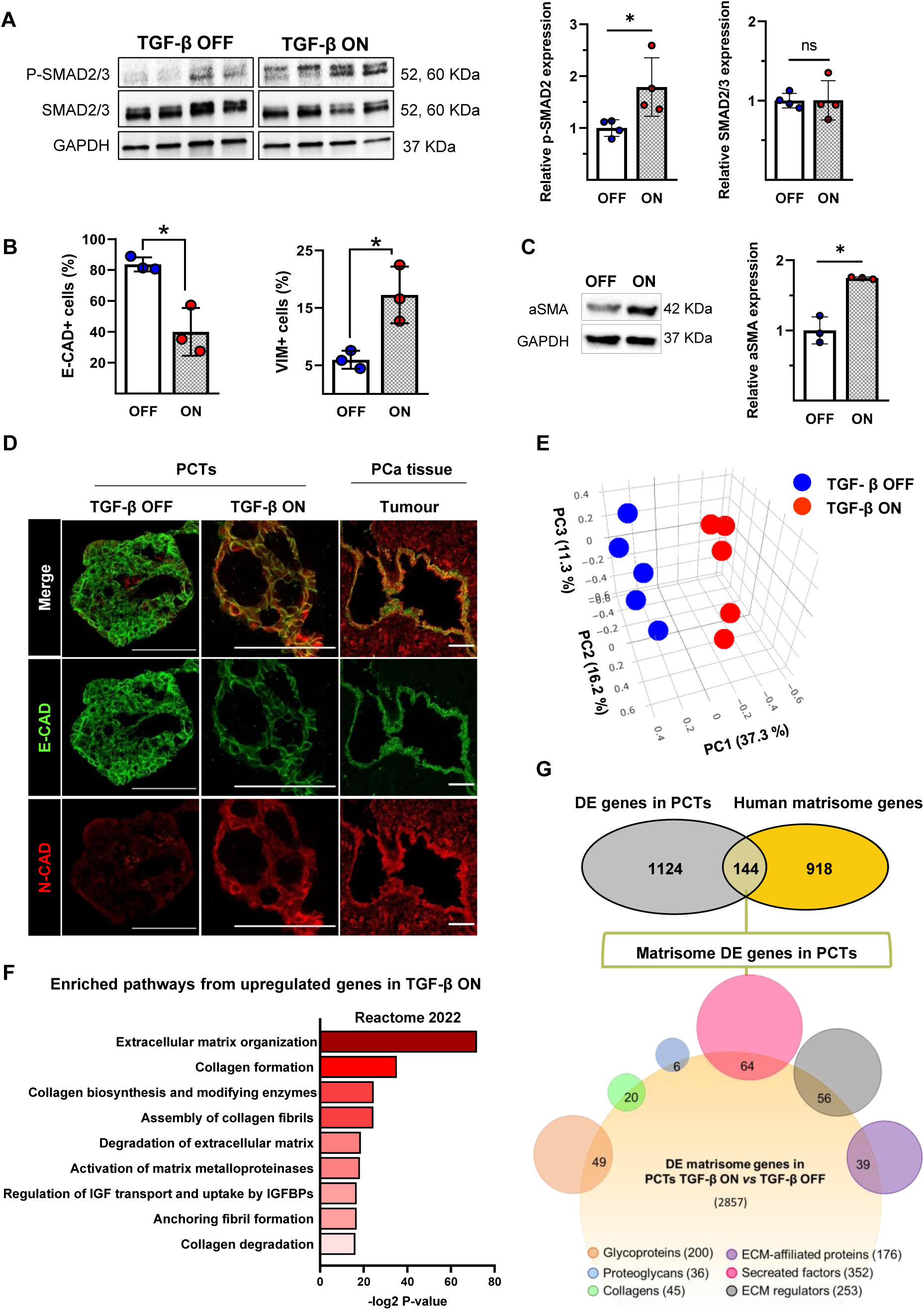
TGF-β activation determines increased expression of EMT markers and ECM pathological remodelling in 3D PCTs. **(A)** SMAD2/3 (N = 4) and p-SMAD2 (N = 4) protein quantification by western blot. Statistical analysis was performed by paired t-test (p < 0.05). **(B)** Flow cytometry analysis of epithelial E-CAD and mesenchymal VIM markers expression in PCTs (N = 3). **(C)** α−SMA (N = 3) protein quantification by western blot analysis. Statistical analysis was performed by paired t-test (p < 0.05). **(D)** Representative confocal images of the PCTs. The tumoroids were fixed with PFA 4% and then embedded in OCT for slicing, prior to immunofluorescence staining. Scale bars: 50 µm. **(E)** Principal component analysis (PCA) representation of the results obtained by RNA-seq of PCTs produced out of 5 prostate cancer patients and treated or not with TGF-β inhibitor (TGF-β OFF or TGF-β ON, respectively). **(F)** Bar plot representation of common enriched Reactome pathways obtained from ENRICHR database, considering the 439 upregulated genes in TGF-β ON (1.5-fold, P adj < 0.05). **(G)** Venn diagram representation of the DE genes found significantly regulated in the PCTs corresponding to the human matrisome, divided by categories.

**Figure 5.**
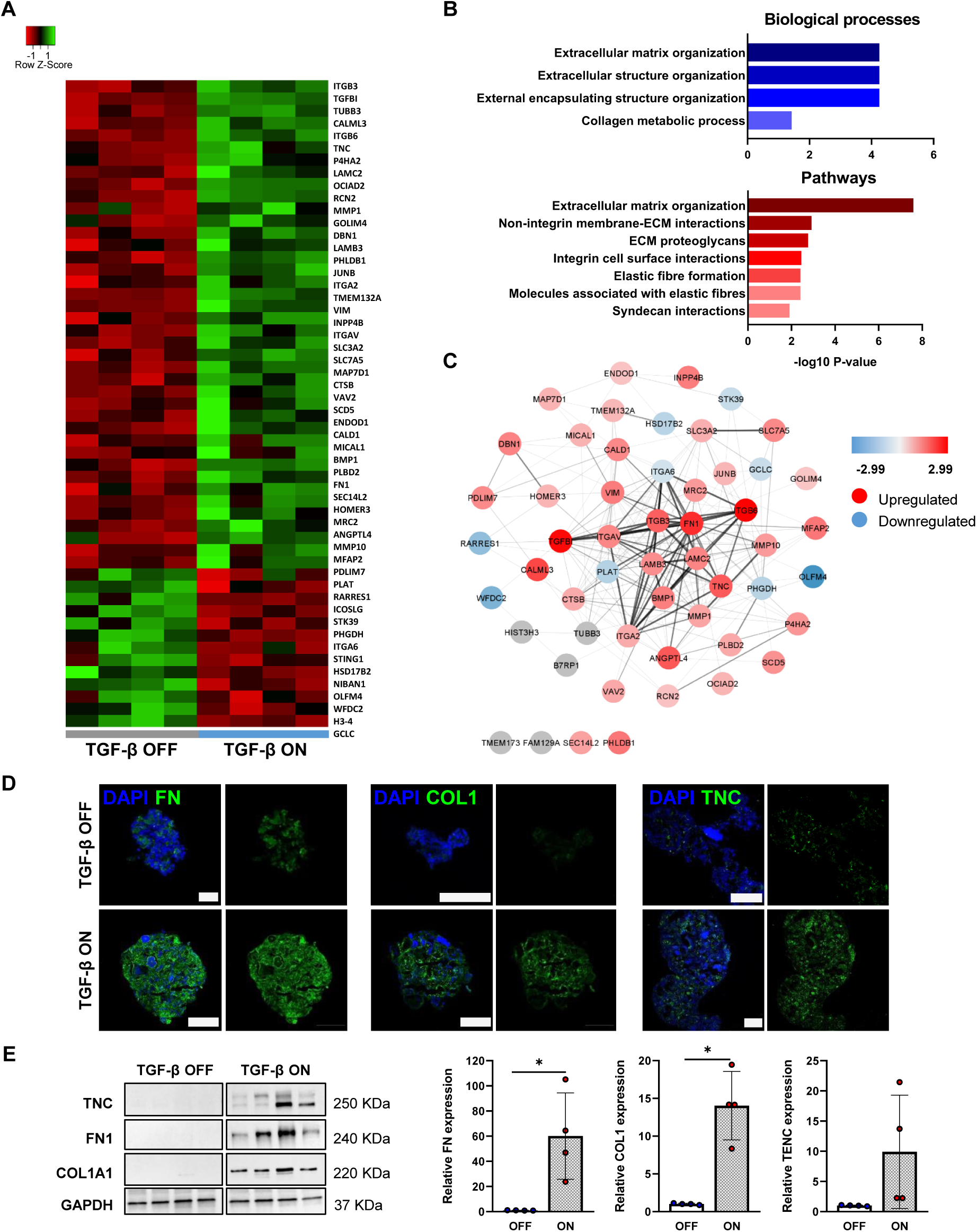
Proteomics analysis unveils TGF-β boosting effect on ECM deposition and remodelling in PCTs. **(A)** Heatmap cluster representation of the 53 proteins found differentially expressed in PCTs treated or not with TGF-β (P adj < 0.05, fold change 0.6). **(B)** Bar plot representation of common enriched biological processes and pathways obtained from ENRICHR database based on the proteins found upregulated in PCTs treated with TGF-β. **(C)** PPI STRING network of the significantly regulated proteins (P adj < 0.05, fold change 0.6) as obtained from Cytoscape App. Nodes are color-coded according to the log2fold change for each protein expression. **(D)** Representative confocal images depicting collagen (COL1), fibronectin (FN) and tenascin (TNC) expression in PCTs stimulated or not with TGF-β. Scale bars: 50 µm. **(E)** Western blot analysis and relative bar plot quantification of COL1, FN and TNC in PCTs stimulated or not with TGF-β. Paired t-test (N = 4).

Next, we examined the transcriptional landscape of the PCTs exposed or not to TGF-β signalling by RNA sequencing (RNA-seq). Principal component analysis (PCA) of the RNA-seq data indicated that the PCTs collected from different patients were particularly sensitive to TGF-β pathway activation, so that PCTs obtained from 5 prostate cancer patients clustered together when stimulated or not with TGF-β (TGF-β ON *vs* TGF-β OFF in Figure 4E). The analysis yielded the differential expression of a total of 1268 genes (P_adj_ < 0.05, |1.5-fold|, Figure S10 and Supplementary Table 2). We focused on the upregulated genes (439) and performed gene ontology (GO) enrichment analysis (Figure 4F). Among the most upregulated genes in the TGF-β ON sample, a significant enrichment for genes involved in ECM organization, formation and degradation could be detected. Consistently, numerous genes encoding for various ECM structural components were deregulated following TGF-β activation (Figure 4G). Venn diagram analysis demonstrated that as many as 144 of the genes found differentially regulated in stimulated PCTs coded for matrisome proteins (Figure 4G and Figure S11).

Among them, we confirmed that TGF-β activation determined the deregulation of numerous members of collagen family (*COL1A1*, *COL7A1*, *COL22A1*, *COL5A1*, *COL6A3*, *COL8A2*, *COL27A1*, *COL4A4*, *COL14A1*), glycoproteins (including *FN1*, *TNC* and *TGFBI*) and proteoglycans (*VCAN*, *SPOCK1*) categories (Figure S11). Additionally, we also identified various genes encoding for ECM regulators (*MMPs*, *ADAMTSs*, *LOXL2* and *BMP1*) and critically involved in ECM degradation and remodelling which were induced after TGF-β stimulation. We corroborated the increased expression of some of the ECM genes in TGF-β-stimulated PCTs by RT-qPCR by using primers for *COL1A1*, *FN1*, *TNC*, *MMPs*, among others (Figure S12).

Overall, these results indicate that TGF-β signalling is critical to the establishment of PCTs in culture. They also indicate that ECM deposition and the acquisition of EMT features in prostate cancer might be caused by TGF-β pathway activation.

### Proteomic profiling confirms TGF-β induced ECM tumorigenic signature of PCTs

Next, we performed quantitative proteomics to understand whether the findings obtained from RNA-seq data and RT-qPCR analysis, which suggests a direct effect of TGF-β on the expression of ECM-related genes in PCTs, were mirrored by changes in the expression of ECM proteins. Mass spectrometry analysis returned a total of 53 proteins differentially expressed following TGF-β pathway modulation (fold change 0.6, P_adj_ < 0.05) with 40 proteins being upregulated and 13 downregulated in TGF-β ON PCTs (Figure 5A). The most regulated proteins were ECM glycoproteins (FN1, TNC) and ECM regulators (MMPs, BMPs). GO enrichment analysis confirmed that TGF-β pathway activation determined a general increase in ECM deposition and organization in PCTs (Figure 5B) which was mirrored by a tightly interconnected network among the ECM components as determined by protein-protein interaction (PPI) analysis using STRING database (Figure 5C). In detail, the interaction between TGFBI, FN1, TNC, ITGV, ITGB3, ITGA6, VIM, MMP10, and BMP1 was detected with high confidence by PPI, suggesting that essential interactions between them are indispensable for ECM remodelling during PCa triggered by TGF-β pathway. Of note, ITGA6 is found downregulated in the sample TGF-β ON which is in accordance with what is observed in the patients’ tissues and is a distinctive characteristic of PCa, according to our results (see Figure 1).

These findings were also corroborated through immunocytochemistry (ICC) and western blot analyses for fibronectin (FN), collagen 1 (COL1) and tenascin C (TNC) (Figure 5D, E) which displayed significantly enhanced expression in PCTs exposed to TGF-β signalling.

### TGF-β-induced alterations are instrumental to prostate cancer cell migration and metastatization

Results previously generated in mouse and cancer cell lines highlighted the role of TGF-β as a potent EMT driver during tumour progression and metastatization for various solid cancers.^1, 31, 32^

To confirm whether this phenomenon could be reproduced in prostate tumoroids, we isolated single cells from PCTs that had been exposed or not to TGF-β and performed a collagen pad contraction assay. We were able to confirm that TGF-β pathway activation – which determined the protein upregulation of key components of the focal adhesions, like ITGA2, ITGB6, ITGB3 and ITGV in PCTs (Figure 6A) – also boosted the ability of prostate tumour-derived cells to remodel the surrounding ECM (Figure 6B). Additionally, the activation of this pathway triggered in PCTs the ability to promote degradation of the surrounding ECM, a necessary feature for tumour cell invasion, as assessed by the degradation of a fluorescent gelatin substrate (Figure 6C and Figure S13). To reinforce these results, we set up an ECM invasion assay, whereby PCTs in which TGF-β signalling pathway was activated or not were embedded in Matrigel^TM^ and checked for the ability to spread out and migrate. As expected, tumour cells migrated out more efficiently from PCTs in which TGF-β pathway had been activated as compared to those in which the pathway had been blocked, the latter remaining in the 3D tumoroid conformation within the Matrigel^TM^ (Figure 6D).

**Figure 6.**
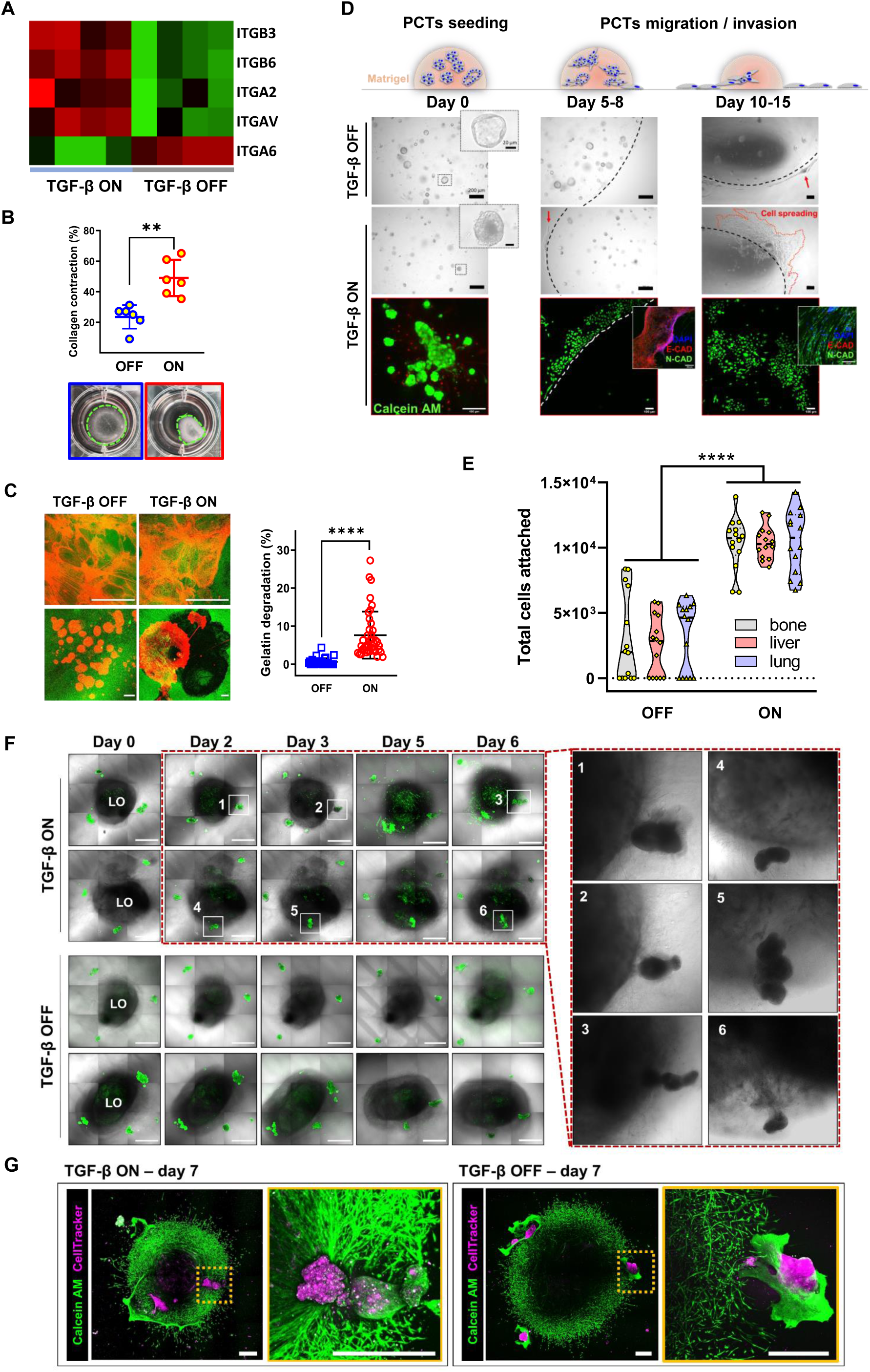
TGF-β drives EMT and enhances the metastatic potential in 3D prostate cancer tumoroids. **(A)** Heatmap cluster representation of the expression of integrins found differentially expressed, by mass spectrometry, in PCTs exposed to TGF-β or not. **(B)** Top: Dot plot representation of the data obtained by contraction assay. The data are shown as percentage of the collagen pads area (N = 6). Bottom: Representative bright field image of the contracted pad (day 7). **(C)** Left: Representative confocal microscope image of gelatin degradation induced by PCTs stimulated or not with TGF-β. Green: gelatin; orange: F-actin. Right: Dot plot representation of the data obtained by fluorescent gelatin degradation assay (N = 4, n ≥ 7). **(D)** Representative bright field and confocal images of PCTs (green), stimulated or not with TGF-β and cultured in Matrigel™. Scale bars for optical images 200 µm and 20 µm for the insets. Scale bars for confocal images: 100 µm. **(E)** Violin plot representation of the data obtained for PCT cell colonization of bone, liver, and lung decellularized matrices (dECMs) in the presence of TGF-β or not (N = 3, n = 5). 2-way ANOVA followed by Sidak’s multiple comparisons test. **(F)** Representative confocal images of the confrontation assay performed using human induced pluripotent stem cell (iPSC)-derived lung organoids (LOs) and PCTs (green). Scale bars: 1000 µm. The insets depict high magnification images of TGF-β-stimulated PCTs invading the LOs. **(G)** Representative confocal microscope images of PCTs (pink) stimulated or not with TGF-β invading hiPSC-LOs (green) at day 7. Scale bars: 500µm.

Finally, we set at clarifying whether TGF-β would affect the efficiency of prostate cancer colonization of the primary sites of metastatization, namely bone, liver and lungs.^33, 34^ For this purpose, we live-stained PCTs which had been stimulated or not with TGF-β with Calcein AM and cultured them onto decellularized matrices obtained from the abovementioned organs. When we measured the number of PCT cells that adhered onto the matrices after 1 hour of incubation, the results revealed that PCTs that had been exposed to TGF-β displayed a significantly higher capacity to attach and spread to the ECM of the three different organs (Figure 6E). Additionally, in order to corroborate this result in a more physiologically relevant model, we set up a 3D confrontation assay in which multiple CellTracker-stained PCTs that had been obtained from two different patients, cultured with or without TGF-β inhibitor, were embedded in Cultrex® hydrogel in the presence of a lung organoid (LO) derived from human induced pluripotent stem cells (hiPSC)^35^. This methodology has been previously adopted to study the migration and invasion properties of tumour spheroids.^36, 37^ PCTs behaviour in the presence of iPSC-derived LOs was monitored over a week of culture by live confocal imaging. Our findings revealed that TGF-β-stimulated PCTs displayed an enhanced ability to migrate and invade the iPSC-derived LOs. In fact, 3 out of 6 PCTs and 5 out of 7 PCTs obtained from two different patients (54 % in total) were able to invade the LOs when the growth factor pathway was active. On the contrary, only 0 out of 6 and 1 out of 6 PCTs from the same patients (8 % in total) succeeded in colonizing the organoid in the absence of TGF-β (Figure 6F and S14). Close-up images taken at day 7 confirmed that TGF-β prompted PCTs integration in the lung organoid stained with Calcein AM (Figure 6G). Altogether, these functional assays confirm, with compelling evidence, that PCTs exposed to TGF-β signalling displayed higher capacity to deform, degrade and cross the surrounding ECM, so that tumour cells can disseminate and more effectively colonize the nearby organs.

### ECM tumorigenic alterations induce EMT in prostate cells

We next set at investigating whether the alterations in tumour ECM determined by TGF-β might be sufficient to affect healthy prostate cell phenotype *per se*, possibly causing them to undergo EMT. To achieve this goal, we decellularized patient primary prostate tumour tissues and used the dECMs as 3D culture substrates to grow prostate cell line PNT2 for 10 days. The successful colonization of the dECMs by PNT2 cells was confirmed by two-photon microscopy 10 days after the cells had been seeded (Figure 7A). Next, we analysed the effect of pathological dECM on PNT2 cell phenotype by staining the cells with an antibody directed against the well-established mesenchymal marker vimentin. To our surprise, two-photon microscopy analysis revealed the presence of the tumour microenvironment was sufficient to induce the expression of vimentin in PNT2 cells (Figure 7B). To confirm this observation, the cells were detached at day 10 from the dECMs and the expression of vimentin was measured by flow cytometry. The analysis confirmed that the tumorigenic ECM promoted the expression of vimentin mesenchymal marker in as much as 77.8 % ± 2.1 compared to 57.3 % ± 4.6 in healthy prostate cells. These striking results were corroborated by using dECMs obtained from two different patients (Figure S15A.

**Figure 7.**
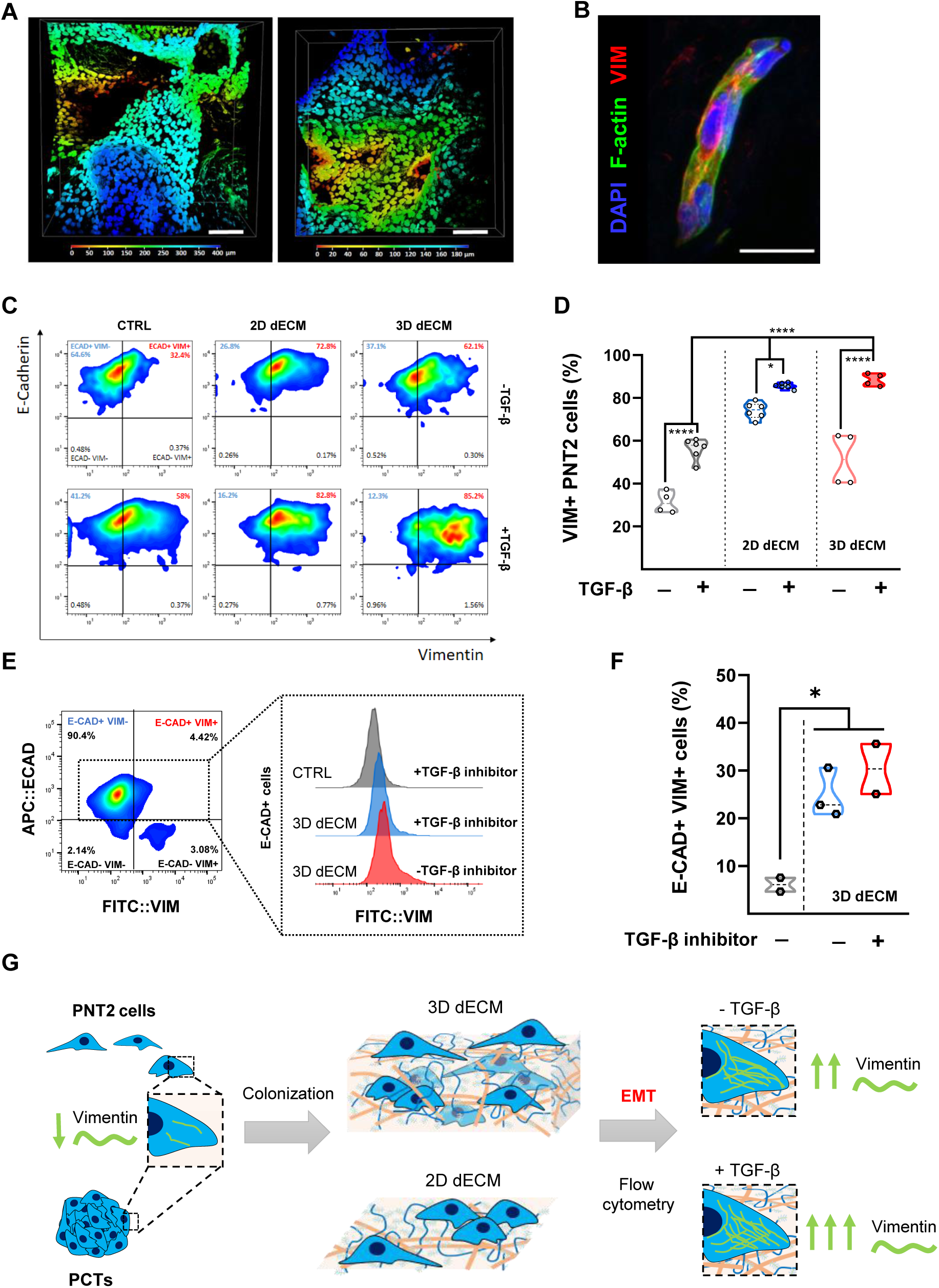
Tumorigenic ECM induces the expression of EMT markers in healthy prostate cells. **(A)** Representative two-photon images of PNT2 cells taken after 10 days of colonization of two tumour dECMs obtained from two different patients. The images represent maximum projections of 3D z-stack images. Cells were stained with DAPI. The colour code refers to the depth to which nuclei are found inside the dECM. Scale bars: 100 µm. **(B)** Representative confocal image of PNT2 cells cultured for 10 days on tumour dECM. The cells are stained for vimentin (VIM, red), F-actin (green) and nuclei are counterstained DAPI (blue). Scale bar: 25 µm. **(C)** Flow cytometry dot plots of vimentin and E-cadherin expression for PNT2 cells, after a week of incubation with 2D and 3D dECMs, in the presence or absence of TGF-β. CTRL cells grown directly on the surface of culture flasks are also shown. **(D)** Vimentin expression quantification based on flow cytometry analysis. Statistical analysis was performed by two-way ANOVA. **(E)** Representative dot plot of PCTs containing E-cadherin (E-CAD), vimentin and dual positive cells. Histogram representation of the epithelial cells from the PCTs expressing vimentin, for control condition and cultured in 3D dECMs in the presence or absence of TGF-β inhibitor **(F)** Vimentin expression quantification based on flow cytometry analysis, for E-CAD-positive cells. Statistical analysis was performed by one-way ANOVA. **(G)** Graphical representation of the experimental design and the results obtained.

Next, we asked whether the changes detected in the chemical composition of the tumour microenvironment were sufficient to trigger the activation of the EMT program in healthy prostate cells. Hence, prostate cancer dECMs were mechanically dissociated to be used as a coating solution for PNT2 cells. Vimentin expression was therefore monitored by flow cytometry and confocal microscopy by using PC3 cancer prostate cell line as a positive control. The results of these analyses demonstrated that the changes in tumour ECM chemical composition were indeed sufficient to boost the expression of vimentin in PNT2 cells (Figure S15B, C).

We then set at investigating whether this effect was directly mediated by the alterations induced by TGF-β in tumour ECM 3D nanostructure and composition. Hence, we repeated the experiment by culturing PNT2 cells, with or without TGF-β stimulation, on 3D or on mechanically dissociated dECMs from other two different patients. After 10 days of culture, we analysed the expression of mesenchymal and epithelial markers vimentin and E-cadherin by flow cytometry. We confirmed that every condition on its own could induce some degree of EMT in PNT2 cells: TGF-β (CTL+TGF-β; 56.2 % ± 4.51), TME chemical composition (2D dECM -TGF-β; 74.1% ± 3.4) and 3D structure (3D dECM -TGF-β; 51.4% ± 10.8). Nevertheless, our analysis demonstrated that the synergistic changes of the 3D ECM structure were boosted by the presence of TGF-β (Figure 7C, D), exerting the most pronounced effect on healthy prostate cells (88.5 % ± 2.5).

These striking results were also confirmed in cells obtained from patient PCTs. Here, E-CAD was used to distinguish the epithelial population of the tumoroids. We observed a significant increment in E-CAD^+^ / VIM^+^ cells after 10 days of culture on 3D tumour dECMs (3D dECM –TGF-β; 24.8 % ± 4.2 *vs* 3D dECM +TGF-β 30.4 % ± 5.3) (Figure 7E, F).

These findings strongly indicate that both the changes in TME chemical composition and its altered 3D structure contribute to promote EMT in healthy prostate cells independently or in combination with TGF-β signalling (Figure 7G).

## Discussion

Amongst the challenges encountered in the fight against cancer, the appropriate representation of the disease *in vitro* is a crucial weakness that needs to be addressed. This is particularly true for prostate cancer, a condition for which reliable and predictive *in vitro* models are currently not available.

Here, we have engaged in the thorough characterization of prostate cancer tissue in a cohort of 80 patients diagnosed with the disease at different stages and undergoing radical prostatectomy.

Besides confirming the disruption of the 3D architecture of the glandular tissue and the loss of basal layer integrity as hallmarks of prostate cancer (PCa) progression, we used scanning electron microscopy (SEM) in combination with two-photon microscopy to scan tumour and non-tumour adjacent tissues and identify reproducible modifications in the distribution and alignment of extracellular matrix (ECM) fibres, which could be monitored locally through the accumulation of collagen and fibronectin. After confirming the perturbation in local tumour microenvironment by computational fluid dynamics (CFD), we moved on to identify signs of *desmoplasia* during prostate cancer progression leading to the occurrence of epithelial-to-mesenchymal transition (EMT) in the patients’ tumour tissue. Although being common to many solid tumours, the occurrence of this phenomenon has been debated in prostate cancer.^9–13, 14, 15^

Next, we set at reproducing these tumorigenic features *in vitro* by establishing a protocol to generate patient derived PCa tumoroids. This appeared to be quite the challenge, since very few examples of prostate cancer organoids derived from the primary tissue had been described thus far, most of the existing protocols relying on metastatic cells as a source and most of all with a rather low efficiency.^26, 27, 38^ We - instead - succeeded in generating tumoroids from the primary tissue of all patients independently of their medical history, which displayed a high variability in terms of shape and cellular composition that reflected the histological differences found in the primary tissue. Although their successful generation has been described for a variety of cancer types^23, 26^, PCa tumoroids (PCTs) were previously shown to promptly lose their tumorigenic signature in culture, with tumour cells being overgrown by benign-like prostate cells of epithelial origin.^28^

This was – in fact – also the case with our protocol, whereby ITGA6-positive cells typical of the glandular basal layer almost completely overgrew N-CAD-positive cells within the first 2 weeks in culture. Hence, we speculated that we might be able to preserve PCa cancerous features *in vitro* by activating TGF-β signalling pathway in PCTs. Despite the dual role of this signalling pathway in cancer progression, its deregulation has been implicated in the aetiopathogenesis of PCa.^17, 29^ in fact, high TGF-β levels were indicated as a risk factor for tumour development and progression, with consistently increased levels found in patients with PCa.^39^ We first confirmed that the treatment of healthy prostate cells with TGF-β increases the expression of the mesenchymal markers (VIM) in these cells, suggesting they might undergo EMT which has been proven important in PCa prognosis since it is correlated with the occurrence of PCa bone metastatization.^40–43^

By modulating TGF-β signalling pathway in patient derived PCTs, we demonstrated its role in ECM accumulation and remodelling in prostate cancer. While revealing that TGF-β dictates the appearance of the desmoplastic phenotype, RNA sequencing in PCTs highlighted that significant changes in the expression of 144 genes belonging to the human matrisome were to be ascribed to TGF-β signalling. These included the regulation of genes which had been previously related to solid tumour progression like collagens, fibronectin, *MMPs*, *BMPs* and integrins.^5^ Interestingly, *COL4A4*, a gene encoding for one of the six subunits of type IV collagen and a major structural component of distinctive basal membranes,^44^ was found downregulated in the samples with TGF-β pathway activated. Likewise, the expression of ITGA6, typical of basal epithelial cells, was found downregulated in the sample TGF-β ON, which mirrors our finding for the patient tissue samples (Figure 1). These results might at least partially account for the loss of basal membrane integrity in the tumour tissues, which is thought to facilitate tumour cell migration and metastatization.^45^ On the contrary, the upregulation of fibrillar type I collagen, one of the main effectors of tumour *desmoplasia*, is linked to tumour cell survival.^4^ The analysis of the transcription landscape of TGF-β-stimulated PCTs also returned the upregulation of the expression of important glycoproteins, *i.e.* fibronectin (*FN1*) and (*TNC*). These alterations were further confirmed in mass spectrometry, western blot and immunohistochemistry. Interestingly, fibronectin accumulation at the tumour invasion site and its interaction with TNC is thought to guide tumour cells outside the primary tumour location.^46, 47^ Likewise, the increment of TNC expression at tumour site was correlated to the incidence of *desmoplasia* and tumour proliferation through the modulation of integrin expression.^48–50^ Consistently, a differential expression profile was found for various integrins in our PCTs, which are believed to contribute to increased cancer cell motility.^51^ In particular, the deregulated expression of ITGA5 and ITGV was shown to be linked to cancer cell migration and invasion using prostate cancer cell line DU145 culture in CAFs-derived dECMs. This phenomenon was associated with the enhanced expression of FN-containing aligned fibres.^46, 52^ Our results, suggest that these alterations occur, at least to a certain extent, in response to TGF-β signalling pathway and come accompanied with the overgrowth of cells expressing mesenchymal markers like vimentin.

Consistent with these observations, our migration experiments indicate that TGF-β promoted the motility of prostate tumour cells. More importantly, prostate cancer cells in which the signalling pathway had been activated displayed enhanced capability to adhere to and colonize dECMs extracted from the three main metastatic sites of prostate cancer, *i.e.* bone, lung and liver. This enhanced invasion capacity seems to be related to their higher ability to remodel and degrade the ECM, which is favoured by the higher expression of MMPs and integrins, as well as the enhanced expression of EMT markers, *i.e.* VIM, N-CAD and α−SMA. RNA-seq and mass spectrometry concurred to identify the overexpression of the numerous MMPs in TGF-β-stimulated PCTs including MMP1, 2, 9, 10, 11, 13,

14 and 16. The enhanced activity of gelatinases MMP2 and MMP9 in these tumoroids was independently confirmed through a gelatin degradation assay. Each of the referred proteolytic enzymes has been described to contribute to cancer progression by enhancing tumour cell migration and invasiveness.^53, 54^

With these results in mind, we finally designed a confrontation assay, whereby PCTs stimulated or not with TGF-β were co-cultured with lung organoids derived from human induced pluripotent stem cells (iPSCs). As expected, this assay corroborated that TGF-β pathway activation promoted prostate cancer cell ability to invade lung organoids.

In conclusion, we have described a protocol to establish patient derived prostate cancer organoids that reliably and reproducibly allows for the *in vitro* translation of given PCa features. By exploiting this resource, we demonstrated that TGF-β signalling critically contributes to the tumorigenic alterations in the ECM to trigger prostate cell phenotype switch with enhanced expression of mesenchymal markers. Our findings highlight the cooperative role of TGF-β signalling and ECM *desmoplasia* in prompting EMT in prostate cells and promoting prostate tumour progression and invasion.

## Acknowledgements

We thank the Genomics and Bioinformatics Core Facilities of CEITEC Masaryk University for their support with data analysis and statistics. For proteomics and phosphoproteomics analysis we thank the Proteomics Core Facility at Sahlgrenska Academy, University of Gothenburg, Sweden. This work was supported by the European Regional Development Fund-Projects MAGNET (Number CZ.02.1.01/0.0/0.0/15_003/0000492) and ENOCH (No. CZ.02.1.01/0.0/0.0/16_019/0000868). Moreover, the project received funding from the European Union’s Horizon 2020 research and innovation programme under the Marie Skłodowska-Curie grant agreement No 872233 and No 860715, from EATRIS-CZ research infrastructure project (MEYS Grant No: LM2018133) within the activity of the Project of the large infrastructures for RDI. Ministry of Health of the Czech Republic - DRO (Institute of Hematology and Blood Transfusion – IHBT, 00023736). Marco Cassani, an iCARE-2 fellow, has received funding from Fondazione per la Ricerca sul Cancro (AIRC) and the European Union’s Horizon 2020 research and innovation programme under the Marie Skłodowska-Curie Grant Agreement No. 800924. This work was supported by Marie Curie H2020-MSCA-IF-2020 MSCA-IF-GF “MecHA-Nano”, Grant Agreement No 101031744. We are thankful to Mgr. Filip Kafka for assisting with the culture of hiPSCs-derived lung organoids, Jana Vašíčková and Jiří Šmíd for helping with samples cryosectioning, and Romana Vlčková and Hana Dulová for technical support. The authors would also like to thank Dr. Ana Rubina Perestrelo for technical assistance with tissue handling and histology procedures.

## Authors’ contributions

S.F. and G.F. conceptualized the study and drafted the manuscript. S.F designed, performed the experiments and analysed the data. J.O.D.L.C., M.C., S.M. and H.D. and assisted in designing, carrying out and interpreting experiments. J.O.D.L.C. and M.C. performed data analysis. S.M. optimized the confrontation assay. A.C., A.L., P.F., G.A. and G.D.M. performed the CFD simulations and analysed the data. H.D. helped handling, processing and characterizing patient tissues and prostate cancer tumoroids culture. V.I. assisted in the analysis of mass spectrometry and RNA-sequencing. V.B. and H.F. generated and characterized hiPSCs-derived lung organoids. P.Fi. collected and provided all patient samples used in this study. All authors contributed to manuscript editing, reviewed and approved the final version.

## Materials and methods

### Isolation and culture of 3D PCTs

Immediately after surgery, the patient tissues were collected into DPBS containing 1 % Penicillin/Streptomycin and 2.5 µg/ml fungizone (Gibco, 15290026), and transported in ice from the hospital to the laboratory. Patient-derived tumoroids were obtained by enzymatic digestion of the minced tissue overnight, using 5 mg/mL collagenase type II (ThermoFisher, 17101015) in DMEM/F12+ medium - supplemented with Glutamax, HEPES and Primocin as shown in Table 1- at 37 °C and 5 % CO_2_. The day after, cells were washed with DMEM/F12+ and further incubated at 37 C for 10 min with Tryple containing 10 µM Y/27632. After, the cells were washed again twice using DMEM/F12+ and passed through subsequently cell strainers of 70 µm and 30 µm. Freshly isolated single cells were seeded at a cell concentration of 2x10^6^ cells/mL in ultra-low attachment plates using DMEM/F12++ supplemented as shown in Table 1. Tumoroids medium was exchanged every 2 days.

**Table 1.** List of supplements used to prepare DMEM/F12++ medium.

| Table 1. List of supplements used to prepare DMEM/F12++ medium. |  |
| --- | --- |
| HEPES (10 mM) | Biowest, HEPES buffer 1M |
| Glutamax (2 mM) | Life technologies, 100X, 35050 |
| Primocin (100 µg/mL) | InvivoGen, ant-pm-2 |
| B-27 | ThermoFisher, 17504044 |
| N-acetylcysteine (1.25 mM) | Sigma Aldrich, A9-165-25G |
| EGF (5 ng/mL) | Peprotech, AF-100-15 |
| FGF-2 (2 ng/mL) | Peprotech, |
| FGF-10 (10 ng/mL) | Peprotech, 100-26 |
| Nicotinamide (10 mM) | Sigma Aldrich, 98-92-0 |
| Prostaglandin E2 (1 μM) | Peprtech, 3632464 |
| Noggin (100 ng/ml) | R&D systems, 6057-NG-100 |
| R-spondin 1 (500 ng/ml) | R&D systems, 4645-RS-250 |
| DHT (1nM) | Selleckchem, S4757 |
| Y-27632 dihydrochloride (10 μM) | Selleckchem, S1049 |

### Culture of 2D cell lines

PNT2 human cell line was purchased from Sigma Aldrich (95012613-1VL) and cultured in RPMI 1640 containing 2 mM Glutamine and 10 % Foetal Bovine Serum (FBS), as suggested by the manufacturer. PC3 human cell line was obtained from Sigma Aldrich (90112714-1VL) and cultured in DMEM medium with 10 % FBS.

### Human induced pluripotent stem cell (hiPSC) culture

Human induced pluripotent stem cells (hiPSCs) (WiCell, DF19-9-7T)^55^ were cultured on Cultrex Extract (Stem Cell Qualified Reduced Growth Factor Basement Membrane, RnD)-coated tissue culture dishes. The mTESR+ medium (STEMCELL) was changed every two days. At 80 % confluency, the cells were passaged using TryPLE (Gibco). For the first 24 hours, the RHO/ROCK pathway inhibitor (10 μM, STEMCELL) was used. The next day, the cells were washed with fresh medium and maintained until the next passaging.

### Differentiation of hiPSCs-derived lung organoids

Lung organoids (LOs) were differentiated from hiPSCs following an adapted protocol already published.^35, 56, 57^ In brief, iPSCs were differentiated into definitive endoderm using the three-step protocol. On the first day, the medium was changed and RPMI 1640 (Gibco) supplemented with Activin A (100 ng/mL, RnD) was added. On the second day, the medium was changed and fresh RPMI 1640 supplemented with Activin A (100 ng/mL) and 0.2 % HyClone-defined FBS (GE Healthcare Bio-Sciences) was added. On the third and fourth days, the medium was changed and fresh RPMI 1640 was supplemented with Activin A (100 ng/mL) and 2 % FBS. After four days of differentiation, the endodermal layer was induced to foregut differentiation with LO basic medium (Advanced DMEM F12 (Gibco) supplemented with N2 supplement (Thermo Fisher Scientific) and B27 supplement (Thermo Fisher Scientific), Glutamax supplement (Thermo Fisher Scientific), penicillin and streptomycin (500 U/mL)) supplemented with Noggin (250 ng/mL, RnD), SB431542 (10 μM, STEMCELL), SAG (1 μM, Tocris) and FGF4 (500 ng/mL, RnD), and CHIR-99021 (2 μM, STEMCELL). After 4 - 5 days, 3D spheroids were formed. The spheroids were collected, embedded in Cultrex RGF Basement Membrane Extract, Type 2 (RnD) drops, and fed with complete LO medium (basic LO medium supplemented with 1 % HyClone-defined FBS and FGF10 (500 ng/mL, RnD). The medium was changed twice a week. The LOs were used for the experiments after 50 days in the culture.

### Histology, immunohistochemistry and immunocytochemistry

Human tissues dissected in cold DPBS and 3D tumoroids were fixed for approx. 8 hours in PFA 4 % and cryoprotected in 30 % sucrose at 4 °C overnight. The following day the tissue was embedded in optimal cutting temperature compound (OCT - Leica Biosystems) and snap frozen in isopentane over dry ice. The frozen specimens were later sectioned into 10 µm tick sections using a cryostat (LEICA CM1950).

For immunohistochemistry, the samples were thawed at room temperature (5-10 minutes) and re-hydrated with DPBS for another 5 minutes. Next, samples were permeabilized with 0.2 % Triton X-100 for 10 minutes at room temperature. After a washing step, the blocking was performed using 2.5 % BSA in DPBS for 30 minutes at room temperature. Primary antibodies were incubated overnight at 4 °C (Table 2) or 2 hours at room temperature and then washed prior to Alexa fluorochrome-conjugated secondary antibodies (1:500, ThermoFisher Scientific) incubation for 1 hour. Controls were performed by incubating with secondary antibodies only.

**Table 2.** List of primary antibodies used.

| Method | Target protein | Dilution | Catalog No. | Manufacturer |
| --- | --- | --- | --- | --- |
| IHC / IF | E-Cadherin | 1:200 | ab1416 | Abcam |
|  | N-Cadherin | 1:50 | sc-7939 | Santa Cruz |
|  | Vimentin | 1:100 | V6630 | Sigma-Aldrich |
|  | Cytokeratin 8 | 1:300 | ab53280 | Abcam |
|  | ITGA-6 (CD49f) | 1:50 | sc-19622 | Santa Cruz |
|  | Collagen I | 1:200 | 600-401-103 | Rockland |
|  | Fibronectin | 1:300 | 28836 | Cell signalling |
|  | Tenascin-C | 1:50 | sc-9871 | Santa Cruz |
| FC | Vimentin | 1:100 | V6630 | Sigma-Aldrich |
|  | E-Cadherin | 1:200 | ab1416 | Abcam |
|  | N-cadherin | 1:50 | sc-7939 | Santa Cruz |
| WB | Collagen 1A1 | 1:500 | 91144 | Cell signalling |
|  | Fibronectin | 1:300 | F0791 | Sigma-Aldrich |
|  | Tenascin C | 1:500 | MA1-26779 | Termofisher |
| | α-SMA | 1:1000 | 19245 | Cell signalling |
|  | p-SMAD 2 | 1:1000 | 3108 | Cell signalling |
|  | SMAD 2/3 | 1:1000 | 8685 | Cell signalling |

LOs were washed with PBS and fixed with 4 % paraformaldehyde (PFA) for 20 minutes at room temperature and washed again three times with PBS. The LO were permeabilized with 0.5 % Triton X-100 in PBS and then washed three times with IF buffer (PBS with 0.2 % Triton X-100 + 0.05 % Tween-20). The samples were blocked with 2.5 % bovine serum albumin (BSA) in IF buffer. The LOs were incubated with anti-Acetyl-α-tubulin (Cell Signaling, clone D20G3), and anti-FOXJ1 (eBioscience, clone 2A5) antibodies in IF buffer + 1 % BSA at 4 °C overnight. The LOs were washed with IF buffer three times and incubated with secondary antibodies in IF buffer + 1 % BSA for one hour at room temperature.

For immunohistochemical analysis of adherent cells, the cells were fixed for 10 minutes in PFA 4 %. Next, samples were permeabilized with 0.2 % Triton X-100 for 10 minutes at room temperature. After a washing step, the blocking was performed using 2.5 % BSA in DPBS for 30 minutes at room temperature. Primary antibodies were incubated overnight at 4 °C (Table 2) or 2 hours at room temperature and then washed prior to Alexa fluorochrome-conjugated secondary antibodies (1:500, ThermoFisher Scientific) incubation. Controls were performed by incubating with secondary antibodies only.

For all samples, the nuclei were counterstained with DAPI (Sigma-Aldrich). The slides were then mounted in anti-fade Mowiol® 4-88 mounting media (Sigma-Aldrich) supplemented with DABCO (Sigma-Aldrich). Images were captured using a Zeiss LSM 780 confocal microscope or a LEICA (TCS SP8). For immunostaining to characterize dECMs, DAPI incubation was used as a DNA-negative control to confirm decellularization efficiency.

Tissues and tumoroids histology was assessed, after cryosectioning, by hematoxylin and eosin (H&E) (Sigma Aldrich) staining. Masson’s Trichrome (Sigma-Aldrich) tissue staining was performed according to the manufacturer’s protocol using celestine blue (0.5 % m/v celestine blue, 5 % m/v ammonium iron-III sulphate dodecahydrate and glycerol in distilled water, Sigma Aldrich) and bouin’s solution (VWR Chemicals). Whole tissue sections were visualized under a slide scanner Zeiss Axio Scan Z1 microscope using the bright-field mode.

### Scanning electron microscopy

Decellularized matrices were fixed for 30 minutes under gentle agitation (12 rpm orbital 154 shaker) in 3 % glutaraldehyde solution in cacodylate buffer (100 mM, Sigma 20840) at room temperature. The samples were washed three times for 10 minutes in cacodylate buffer, dehydrated via an ethanol series (10-100 %) for 10 minutes per incubation and then stored at 4 °C until transfer for drying and coating. Dried samples were fixed on holders with carbon tape, sputter-coated with platinum (JFC-1300 autofine coater - JEOL, 40 mA, 15 s) and visualized under a JCM-6000 benchwork scanning electron microscope (Jeol) under high-vacuum, using a SEI detector PC-standard, with accelerating voltage to 15 kV at x2000. Image analysis is detailed in Online Table II.

### Multiphoton microscopy

Second Harmonic Generation (SHG) was obtained using a Carl Zeiss LSM 7 MP Multiphoton Microscope (Ti:Sapphire Laser - Chameleon 690-1080 nm) and a Plan-Apochromat 20x water dipping objective, with 15-35 % laser power, λex = 750-800 nm.

Two-photon-excited fluorescence (TPEF) modality was used for the 3D imaging of PNT2 cells colonization of tumour dECMs, using LP555+BP570-610IR+nm, LP490+BP500-550IR+nm and mirror+SP485 nm filter cubes, resulting in dual-colour visualization of Alexa Fluor 555, Alexa Fluor 488 and DAPI.

### Assessment of samples anisotropy

The methodology consists in assessing anisotropy by first evaluating the *average effective diffusivity tensor* (Eq. 1),^25^ which represents the overall results of the deviation of the diffusional paths with respect to the ideal straight ones (corresponding to the unconstrained free diffusivity). After obtaining the diffusivity tensor, we calculate its eigenvalues along with the corresponding eigenvectors, from which the (diagonal) tortuosity and connectivity tensors are obtained and in turn, from them, the anisotropy factors.^25^

### Methodology

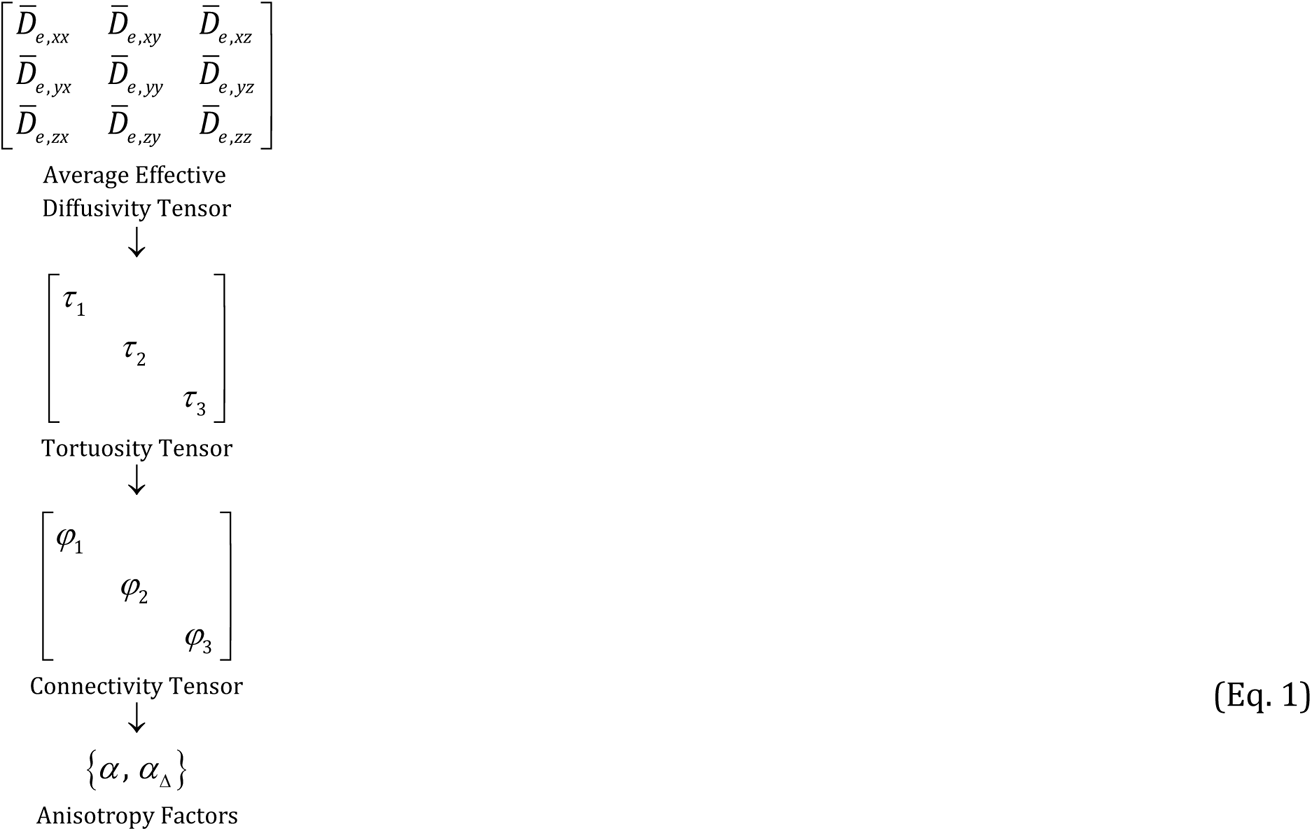

In this work, besides the *anisotropy factor α* introduced by Basser and Pierpaoli (1996), and calculated as reported in Eq. 2,^25^ we additionally introduce and use here the *deviatoric anisotropy factor α*_Δ_ as an alternative way of assessing the anisotropy of the considered tissues. This factor is calculated in analogy to Eq. 2 from the eigenvalues of the deviatoric tortuosity tensor, splitting the isotropic tensor of the minimum tortuosity from the overall tortuosity tensor (Eq. 3).

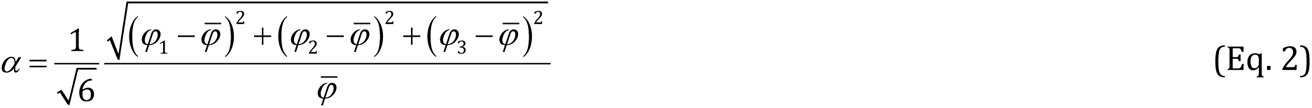

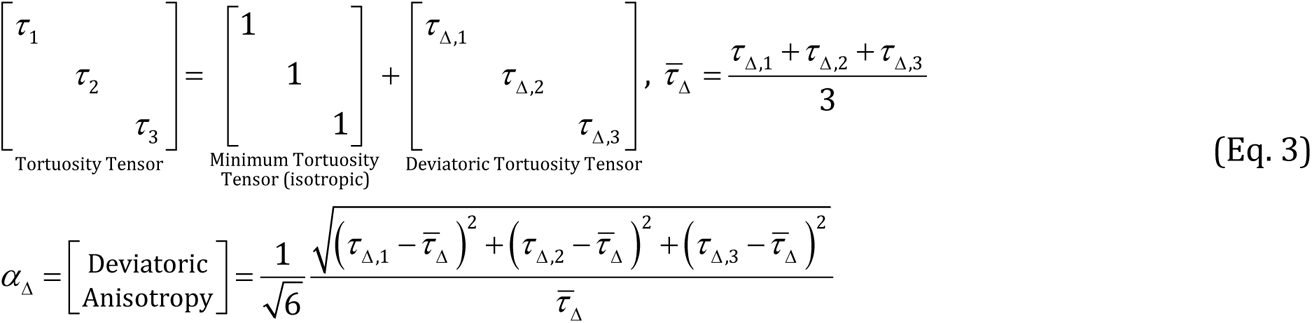

The difference between the two anisotropy factors lies in their different reference values for tortuosity – the unity for the anisotropy factor and zero for the deviatoric one –, which makes the latter more sensitive to the structural anisotropy than the former.

To give some simulation details, the diffusion field is evaluated by solving the 3D Laplace equation, corresponding to the conservation of mass of a species using the Fickian pure-diffusion equations with local uniform free diffusivity as the constitutive equations for flux. The governing equations are discretised with the Finite-Volumes Methods (FVM), which ensures the conservation of mass within domain. The numerical model is implemented in the environment of the open-source software OpenFOAM^®^ through C++ libraries.

The computational domain is discretised by an unstructured mesh, where the dependent variables are stored at the center of each cell with a co-located arrangement. As regards the unstructured mesh-refinement study, various numerical tests were performed before reaching the final mesh configuration using an increasing number of cells in all the directions to ensure the mesh-independency of results.

To obtain the complete average diffusivity tensor, three different simulations are necessary, imposing each time a non-null concentration difference along the *x*-, *y*- and *z*-direction of the cell, respectively. This is done by setting uniform low- and high-concentration values on the two external surfaces normal to the considered flux direction. A null concentration gradient (*i.e.,* null flux) is set on the remaining external surfaces.^25^

### RNA extraction

RNA extraction from tumoroids was done following the manufacturer protocol using the High Pure RNA Isolation Kit (Roche).

### RNA-sequencing and differential expression (DE) analysis

#### Sequencing library preparation

100 ng/µl of total RNA per sample was used as an input for library preparation using QuantSeq FWD 3’mRNA Library Prep Kit (Lexogen). Briefly, RNA was transcribed into cDNA using oligodT primer (FS1) at half volume compared to the manufacturer’s instructions to minimize the off-target products. Following the first strand synthesis, RNA removal and second strand synthesis were performed using the UMI Second Strand Synthesis Mix (USS) containing Unique Molecular Identifiers (UMIs) that allow detection and removal of PCR duplicates. Finally, sequencing libraries were created by PCR with i5 Unique Dual Indexing Add-on Kit for Illumina (Lexogen). The quality and quantity of libraries were determined using Fragment Analyzer by DNF-474 High Sensitivity NGS Fragment Analysis Kit (Agilent Technologies) and QuantiFluor dsDNA System (Promega). Final library pool was sequenced on Illumina NextSeq 500 with 75 bp single-ends, producing about 10 million reads per library.

#### Data analysis

High-throughput RNA-Seq data were prepared using Lexogen QuantSeq 3’ mRNA-Seq Library Prep Kit FWD for Illumina with polyA selection and sequenced on Illumina NextSeq 500 sequencer (run length 1x75 nt). Bcl files were converted to Fastq format using bcl2fastq v. 2.20.0.422 Illumina software for basecalling. 6-nt long UMIs were extracted and subsequently used for deduplication of aligned reads by UMI-tools v. 1.1.1.^58^ As a next step 6-nt long barcode sequence related to Lexogen QuantSeq Library Prep Kit were trimmed using seqtk 1.3-r106.^59^ Quality check of raw single-end fastq reads was carried out by FastQC v0.11.9.^60^ The adapters and quality trimming of raw fastq reads was performed using Trimmomatic v0.36^61^ with settings CROP:250 LEADING:3 TRAILING:3 SLIDINGWINDOW:4:5 MINLEN:35 and adaptor sequence ILLUMINACLIP:AGATCGGAAGAGCACACGTC. Trimmed RNA-Seq reads were mapped against the human genome (hg38) and Ensembl GRCh38 v.94 annotation using STAR v2.7.3a^62^ as splice-aware short read aligner and default parameters except -- outFilterMismatchNoverLmax 0.1 and --twopassMode Basic. Quality control after alignment concerning the number and percentage of uniquely and multi-mapped reads, rRNA contamination, mapped regions, read coverage distribution, strand specificity, gene biotypes and PCR duplication was performed using several tools namely RSeQC v2.6.2,^63^ Picard toolkit v2.18.27^64^ and Qualimap v.2.2.2^65^ and BioBloom tools v 2.3.4-6-g433f.^66^ The differential gene expression analysis was calculated based on the gene counts produced using featureCounts tool v1.6.3^67^ with settings -s 2 -T 10 -F GTF -Q 0 -d 1 -D 25000 and using Bioconductor package DESeq2 v1.20.0.^68^ Data generated by DESeq2 with independent filtering were selected for the differential gene expression analysis to avoid potential false positive results. Genes were considered as differentially expressed based on a cut-off of adjusted p- value <= 0.05 and log2(fold-change) ≥1.5 or ≤-1.5. PCA and volcano plot were produced using Biojupies web tool. Clustered heatmaps were generated from differentially expressed genes using Heatmapper web tool. Bar plot representation of common enriched pathways was obtained from ENRICHR database, considering the 439 upregulated genes in TGF-β ON.

### Mass spectrometry

#### Sample preparation and digestion

Samples were homogenized on a FastPrep®-24 instrument (MP Biomedicals, OH, USA) for 5 repeated 40-second cycles at 6.5 m/s in lysis buffer containing 2 % sodium dodecyl sulfate (SDS), 50 mM triethylammonium bicarbonate (TEAB) and X1 Pierce Phosphatase inhibitor (A32957, Thermo Fischer Scientific, Waltham, MA, USA). Lysed samples were centrifuged at 8000 g for 20 minutes and the supernatants were transferred to clean tubes. Protein concentrations were determined using Pierce™ BCA Protein Assay Kit (Thermo Fischer Scientific) and the Benchmark™ Plus microplate reader (Bio-Rad Laboratories, Hercules, CA, USA) with bovine serum albumin (BSA) solutions as standards.

Aliquots containing 30 µg of total protein from each sample were incubated at 60 °C for 30 minutes in the lysis buffer with DL-dithiothreitol (DTT) at 100 mM final concentration. The reduced samples were processed using the modified filter-aided sample preparation (FASP) method.^69^ Briefly, the samples were transferred to 30 kDa Microcon Centrifugal Filter Units (catalogue no. MRCF0R030, Merck), washed repeatedly with 8 M urea and once with digestion buffer prior (0.5 % sodium deoxycholate (SDC) in 50 mM TEAB) to alkylation with 10 mM methyl methanethiosulfonate in digestion buffer for 30 minutes. Digestion was performed in digestion buffer by addition of Pierce MS grade Trypsin (Thermo Fisher Scientific), in an enzyme to protein ratio of 1:100 at 37 °C overnight. An additional portion of trypsin was added and incubated for 4 hours. Peptides were collected by centrifugation.

Isobaric labelling was performed using Tandem Mass Tag (TMTpro 16plex) reagents (Thermo Fischer Scientific) according to the manufacturer’s instructions. The labelled samples were combined into one pooled sample, concentrated using vacuum centrifugation, and SDC was removed by acidification with 10 % TFA and subsequent centrifugation. The labelled pooled samples were treated with Pierce peptide desalting spin columns (Thermo Fischer Scientific) according to the manufacturer’s instructions The total proteome sample was pre-fractionated using basic reversed-phase chromatography (bRP-LC) with a Dionex Ultimate 3000 UPLC system (Thermo Fischer Scientific) on a reversed-phase Xbridge BEH C18 column (3.5 μm, 3.0x150 mm, Waters Corporation), via a linear gradient from 3 % to 8 % solvent B over 1 minute, 8 % to 40 % solvent B over 25 minutes, 40 % to 55 % solvent B over 9 minutes, followed by an increase to 100 % B over 5 minutes. Solvent A was 10 mM ammonium formate buffer at pH 10.00 and solvent B was 90 % acetonitrile, 10 % 10 mM ammonium formate at pH 10.00. The fractions of the whole proteomics fractions were concatenated into 19 fractions dried and reconstituted in 3 % acetonitrile, 0.1 % formic acid.

#### nLC-MS/MS

Each fraction was analyzed on an Orbitrap Fusion™ Tribrid™ mass spectrometer interfaced with Easy- nLC1200 liquid chromatography system (Thermo Fisher Scientific). Peptides were trapped on an Acclaim Pepmap 100 C18 trap column (100 μm x 2 cm, particle size 5 μm, Thermo Fischer Scientific) and separated on an in-house packed analytical column (75 μm x 40 cm, particle size 3 μm, Reprosil-Pur C18, Dr. Maisch). The fractions were separated using a linear gradient from 5 % to 12 % B over 5 min, 12 % to 35 % over 70 minutes followed by an increase to 100 % B for 5 minutes, and 100 % B for 10 minutes at a flow of 300 nL/min. Solvent A was 0.2 % formic acid and solvent B was 80 % acetonitrile, 0.2 % formic acid.

MS parameters for the whole proteome; MS scans were performed at 120 000 resolution, m/z range 375-1500. MS/MS analysis was performed in a data-dependent, with top speed cycle of 3 seconds for the most intense doubly or multiply charged precursor ions. Precursor ions were isolated in the quadrupole with a 0.7 m/z isolation window, with dynamic exclusion set to 10 ppm and duration of 45 seconds. Isolated precursor ions were subjected to collision induced dissociation (CID) at 30 collision energy with a maximum injection time of 50 ms. Produced MS2 fragment ions were detected in the ion trap followed by multinotch (simultaneous) isolation of the top 10 most abundant fragment ions for further fragmentation (MS3) by higher-energy collision dissociation (HCD) at 55 % and detection in the Orbitrap at 50 000 resolutions, m/z range 100-500.

#### Proteomic Data Analysis

The data files of the fractions from each set and study were merged for identification and relative quantification using Proteome Discoverer version 2.4 (Thermo Fisher Scientific). Identification was performed using Mascot version 2.5.1 (Matrix Science) as a search engine by matching against the human database of SwissProt. The precursor mass tolerance was set to 5 ppm and fragment mass tolerance to 0.6 Da (whole proteome). Tryptic peptides were accepted with zero missed cleavages, variable modifications of methionine oxidation and fixed cysteine alkylation, TMTpro-label modifications of N-terminal and lysine were selected. Percolator was used for the validation of identified proteins.

Quantification was performed in Proteome Discoverer 2.4. TMTpro reporter ions were identified with 3 mmu mass tolerance in the MS3 HCD spectra. Only the quantitative results for the unique peptide sequences and the average S/N above 10 were taken into account for the protein quantification and for the whole proteome with the minimum SPS match % of 65. Peptides and proteins were filtered for high confidence.

The quantitative proteomic analysis was run through a KNIME environment workflow. Different normalization methods were tested (quantile, median and loess) and loess method was used. Differential expression analysis was then run using R limma package.

#### ELISA

Elisa kits for the quantification of PSA (DY1344) and TGF-β (DY1344) were purchased from R&D systems. All procedures and measurements were performed as suggested by the manufacturers. Cell culture supernatants were collected from the tumoroids culture, centrifuged at 13000 rpm for 10 minutes and stored at -80 °C until further use.

#### Total collagen quantification

The total amount of collagen, regardless of its type or maturation, was quantified by Total Collagen Assay (QuickZyme Biosciences) accordingly to the manufacturer’s instructions. Here, tissue samples from tumour and adjacent regions were sliced in approximately 2x2x2 mm explants and used to quantify the collagen present in native tissue (cells plus ECM). The hydroxyproline absorbance of the standards and samples was read (570 nm) in duplicate wells using a Multiskan™ GO Spectrophotometer (ThermoFisher Scientific). The collagen content was normalized to the total amount of protein quantified by BCA Protein Assay Kit (ThermoFisher). Albumin serum bovine standards for calibration; the absorbance was read on a Multiskan™ GO Spectrophotometer for 562 nm.

#### Western blotting

Tumoroids were spin down and the pellet was treated with RIPA buffer (Merk Millipore) with protease and phosphatase inhibitor cocktails (1 % v/v, Sigma-Aldrich), by vortexing every 10 minutes for 5 cycles intercalating with 10 minutes on ice. The cell lysate was centrifuged at 13,000x g for 10 minutes at 4 °C and the protein-containing supernatant was collected. Protein levels were quantified using a Pierce BCA Protein Assay Kit (ThermoFisher) with albumin serum bovine standards for calibration; the absorbance (562 nm) was read on a Multiskan™ GO Spectrophotometer. Pre-boiled protein extracts (5-10 μg) were loaded on 10 % or 4–20 % Mini-PROTEANR TGX™ Precast Protein Gels and separated at 100 V in Tris/Glycine/SDS buffer in Mini-Protean Tetra System (Bio-Rad). The proteins were transferred to a polyvinylidene difluoride membrane (PVDF, Bio-Rad) using the Trans-Blot Turbo transfer system (Bio-Rad). PVDF membranes were blocked with 5 % skim milk or 5 % BSA in TBST for 1 hour, followed by primary antibody incubation in 5 % BSA in TBST with gentle rotation overnight at 4 °C. After several washes in TBST, the PVDF membranes were incubated with the appropriate secondary HRP-conjugated antibody (Sigma-Aldrich) for 1 hour at room temperature. After the membranes were washed 3x during 5 minutes in TBST and the chemiluminescence was detected using Clarity™ Western ECL Substrate (Bio-Rad) and ChemiDoc MP Imaging System (Bio-Rad). Quantification was done using Image Lab 6.1. All antibodies were used according to the manufacturer’s instructions. Their specificity was previously confirmed in western blot analysis by identifying a single or prominent band at the appropriate molecular weight. Antibodies and their working concentrations are detailed in Online Table II.

#### Collagen contraction assay

Collagen contractility was evaluated by collagen pad contraction assay, as described in the literature.^70^ Briefly, cells (4x10^5^ cells/mL) were re-suspended in collagen I solution, rat tail (1 mg/mL final concentration; Gibco A10483) and then 500 µL of the cell mixture was transferred to a 24-well plate for gelification in the presence of NaOH. The gel was monitored overtime for size reduction.

#### Gelatin degradation assay

Cells’ ability to degrade the ECM was evaluated based on a fluorescent gelatin degradation assay. For that, the wells of ibidi µ-slide angiogenesis uncoated plates were first coated with poly/L/Lysine (PLL) (50 µg/mL) at room temperature for 20 minutes. After 3 washes with DPBS, crosslinking was done with glutaraldehyde 0.5 % in DPBS for 15 minutes at room temperature. Again, 3 washes with DPBS were done before coating the wells with fluorescent gelatin 1 mg/mL for 10 minutes in the dark. The gelatin coating solution was prepared by mixing gelatin solution type B (G1393, Sigma Aldrich) with gelatin from pig skin, fluorescein-conjugate (G13187, Thermofisher), at a ratio of 4:1 (v/v) in DPBS. After coating, 3 other washes were performed followed by incubation with 70 % ethanol at room temperature for 30 minutes in the dark. The ethanol was washed out by DPBS (3x) and then incubation with media containing FBS was done to quench free aldehydes for 30 minutes. After, 50 µL of cell suspension containing 1500 cells were seeded per well. After 48 hours the cells were fixed with PFA 4 % for 10 minutes, stained for F-actin 555 and analysed at the confocal microscope.

#### Cell adhesion to different ECMs

Cell adhesion to bone (sigma CC170), liver (sigma CC174) and lung (sigma CC175) ECMs were analysed following the manufacturer’s instructions with minor alterations. Briefly, each well (from a 96 multi-well plate) was coated with 40 µL of ECM solution (1.5 mg/mL) for 45 minutes at 37 °C and 5 % CO_2_. Meanwhile, patient prostate cancer cells were treated with Tryple in order to obtain single cells and stained with Calcein AM. After the gelation was complete, 20 000 cells were added per well and left to incubate for 1 hour. Then, the cell suspension was removed, an additional step of washing with PBS 1X was done to ensure the removal of all cells not attached to the ECM, and the number of cells attached was estimated by using a fluorimeter to quantify Calcein AM signal. The total amount of cells was quantified based on the calibration curve performed using Calcein AM stained cells at different concentrations (serial dilutions: 20 000 – 625 cells per well).

#### PTN2 cells and tumoroids adhesion to tumour dECMs

After decellularization, tumour dECMs were used as previously reported by our group.^24^ The dECMs were either used directly as a 3D scaffold or mechanically homogenized in 400 µL of sterile PBS supplemented with 1 % penicillin/streptomycin and disrupted by zirconium beads (3.0 mm diameter beads in 2 mL prefilled tubes, Benchmark Scientific) using a microtube homogenizer (BeadBug TM D1030-E, Benchmark Scientific). Samples were homogenized up to full dissociation in cycles of 1 minute at 3000 rpm and 2 minutes on ice, to prevent overheating. Protein content was quantified with Pierce BCA Protein Assay Kit (ThermoFisher) according to manufacturer’s instructions. Plates were coated with 5 µg/mL of homogenized ECM in PBS 1 % Penicilin/Streptomycin solution (100 µL/cm^2^) overnight at 37 °C. ECM solution was removed immediately before cell seeding at a concentration of 5 000 cells/cm^2^.

#### Confrontation assay between hiPSCs-derived LOs and PCTs

The hiPSCs-derived LOs were embedded in Cultrex and before gelation was completed, CellTracker-stained tumoroids were placed in close proximity to the LOs, using a micropipette. Then, the co-culture was followed by live confocal microscopy to assess the migration and invasion of the tumoroids towards the LOs. After a week, representative co-cultures for each condition were stained for Calcein AM and imaged at the confocal microscope, prior to fixation in PFA 4 %. Next, the samples were transferred to sucrose 15 % overnight at 4 °C and finally included in OCT for cryosectioning.

#### PCTs chemotherapeutic treatment

Random allocation of the tumoroids was done for performing the test in order to have different sized tumoroids fairly distributed between control and treated samples. All drug solutions were prepared in DMEM/F12++ medium. PCTs were treated with the drug dosages specified in the main text and the viability was quantified by luminescent CellTiter-Glo® 3D assay (G9681, Promega). Representative images of PCTs viability were obtained at the confocal microscope using LIVE/DEAD™ Viability/Cytotoxicity Kit (L3224, Thermofisher Scientific) which stains live cells with calcein AM in green and dead cells with red ethidium homodimer 1.

