## Supporting information for "TGF-β induces matrisome pathological alterations and EMT in patient-derived prostate cancer tumoroids"

**Supplementary Table 1. Enrolled patients summary (N=80)**

| **Gleason score** | **Number of samples (%)** |
|---|---|
| 3+3 | 26 (32.5%) |
| 3+4 | 28 (35.0%) |
| 4+3 | 12(15.0%) |
| 4+4 | 6 (7.50%) |
| 5+4 | 2 (2.50%) |
| Not available | 6 (7.50%) |
| **Age (years)** | 50 - 80 |
| **PSA (ng/mL)** | 1.9 - 34 |
| **BMI** | 28.6 ± 4.6 |

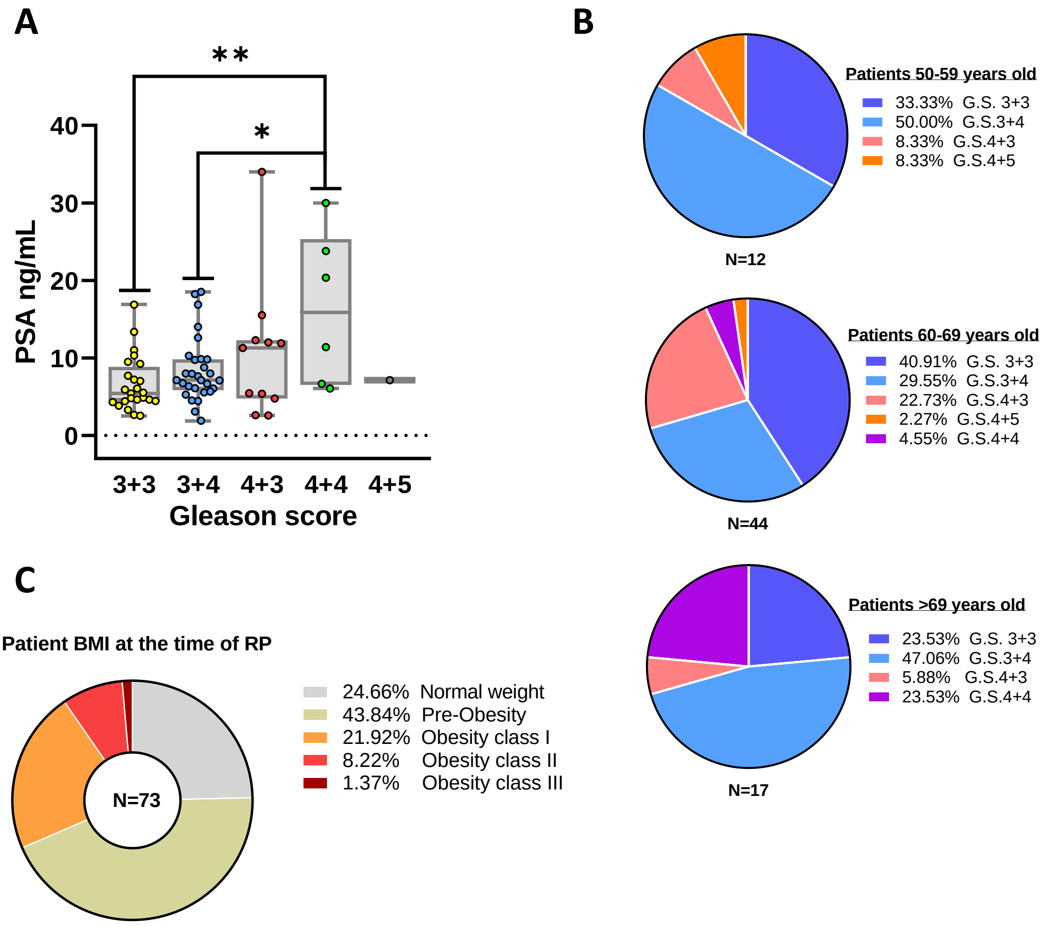


**Figure S1 –** Summary of patients' pathophysiological features. (A) Bar plot representation of the results of prostate specific antigen (PSA) blood level analysis for each patient, grouped by Gleason score. (B) Pie chart representation of patients' age distribution, grouped by Gleason score. (C) Pie chart representation of the blood levels of patients' physical characteristics

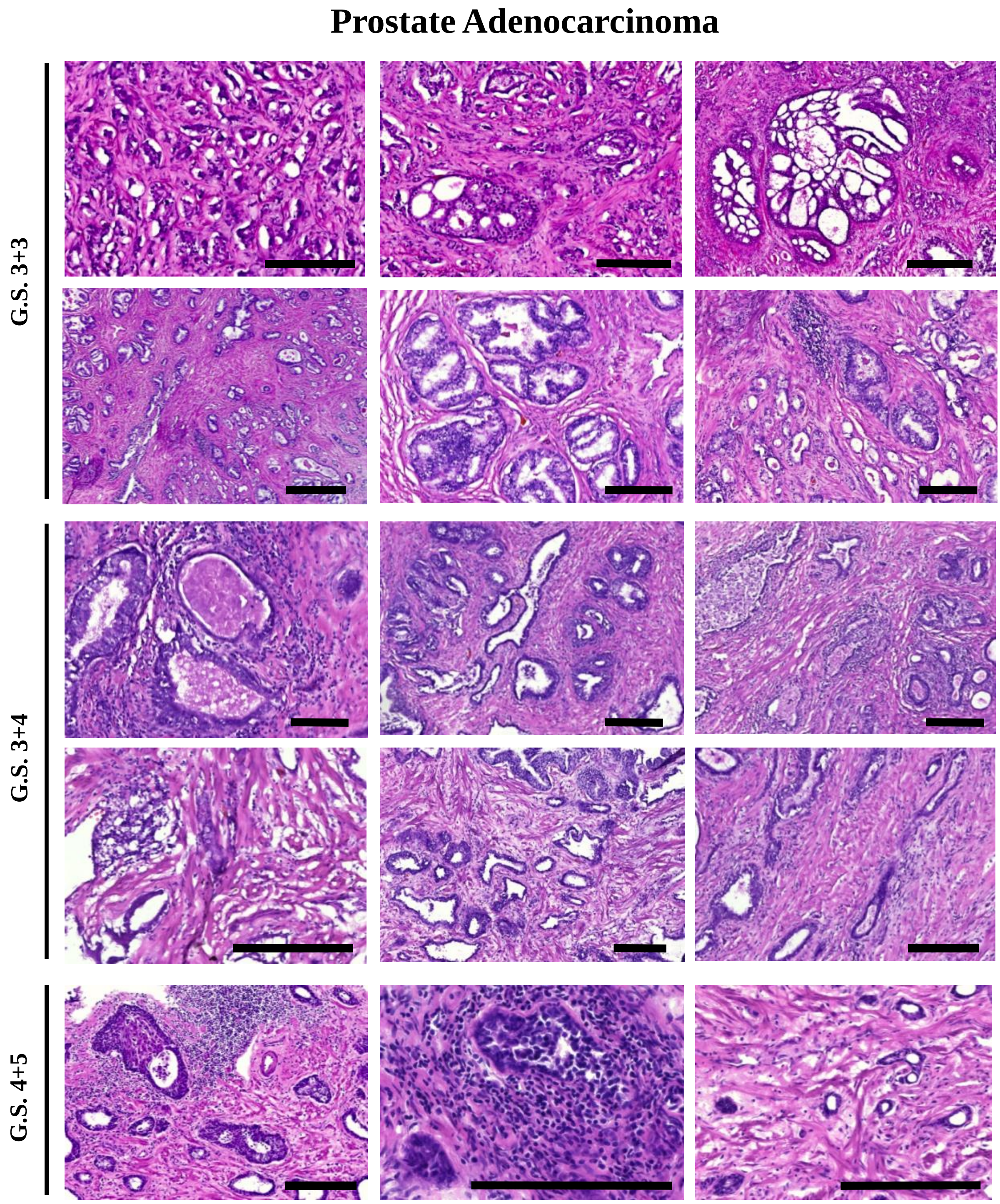


**Figure S2** – Representative H&E histological samples of prostate tumour samples with different Gleason Scores (G.S.). Scale bars: 200 µm.

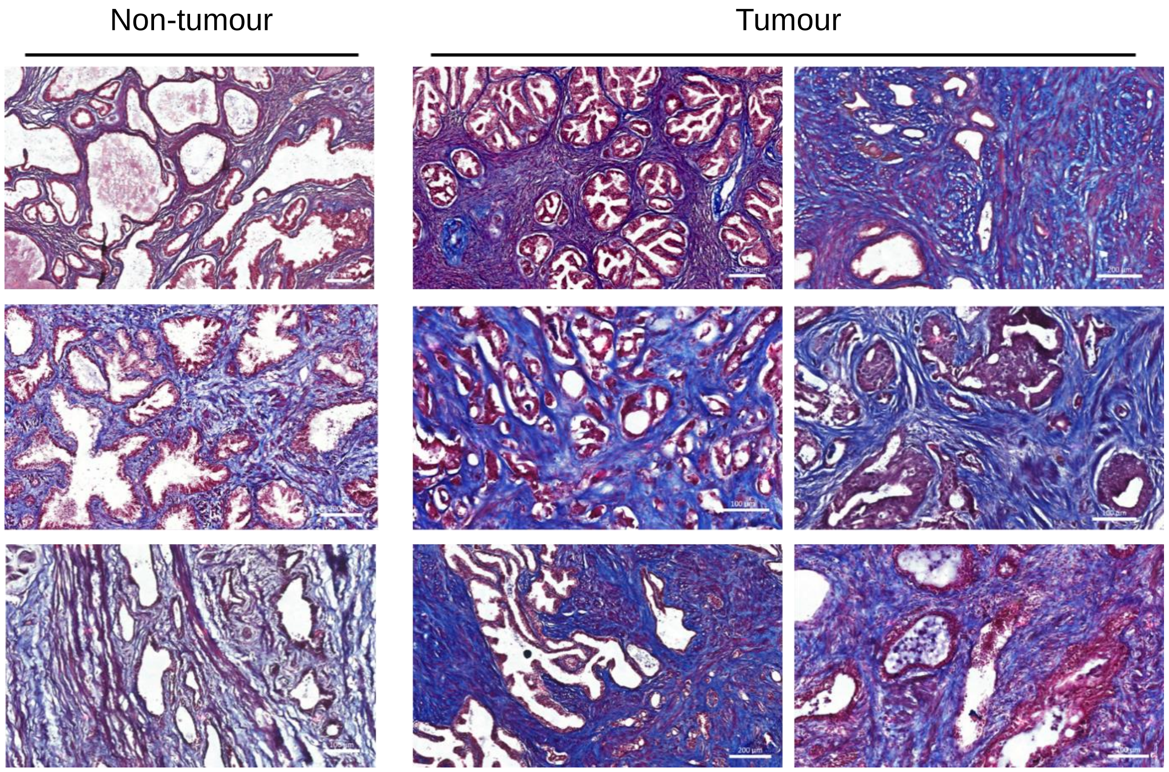


**Figure S3** – Representative Masson's trichrome staining of prostate tumour samples with different Gleason Scores (G.S.) in comparison to their adjacent non tumour tissue.

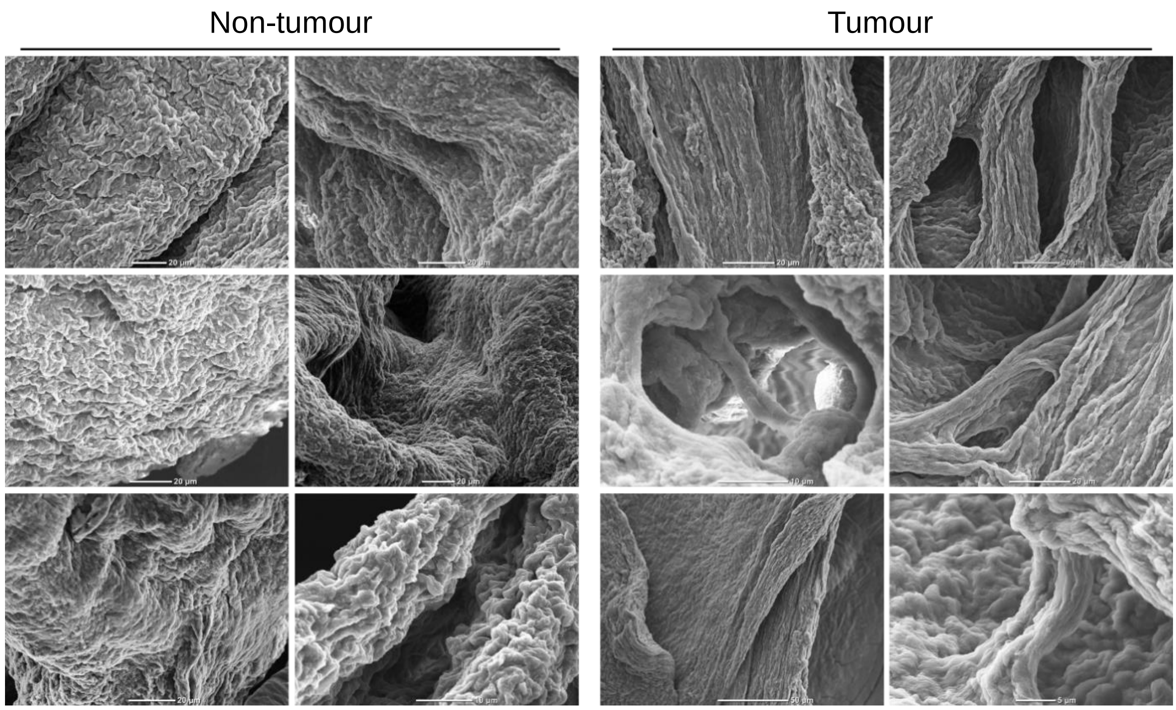


**Figure S4** – Representative SEM images of the surface topography of decellularized extracellular matrices (dECMs) obtained from tumour and adjacent non-tumour prostate tissue obtained from different patients.

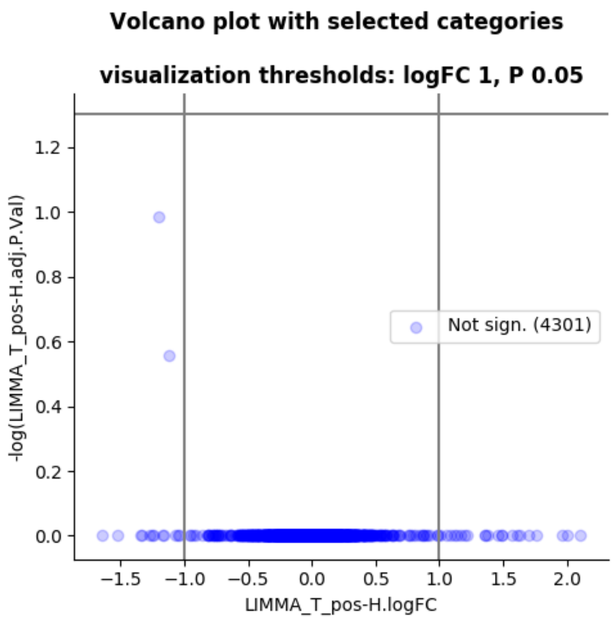


**Figure S5** – Volcano plot representation of up- and downregulated proteins in PCTs (1-fold up or down, P adj < 0.05).

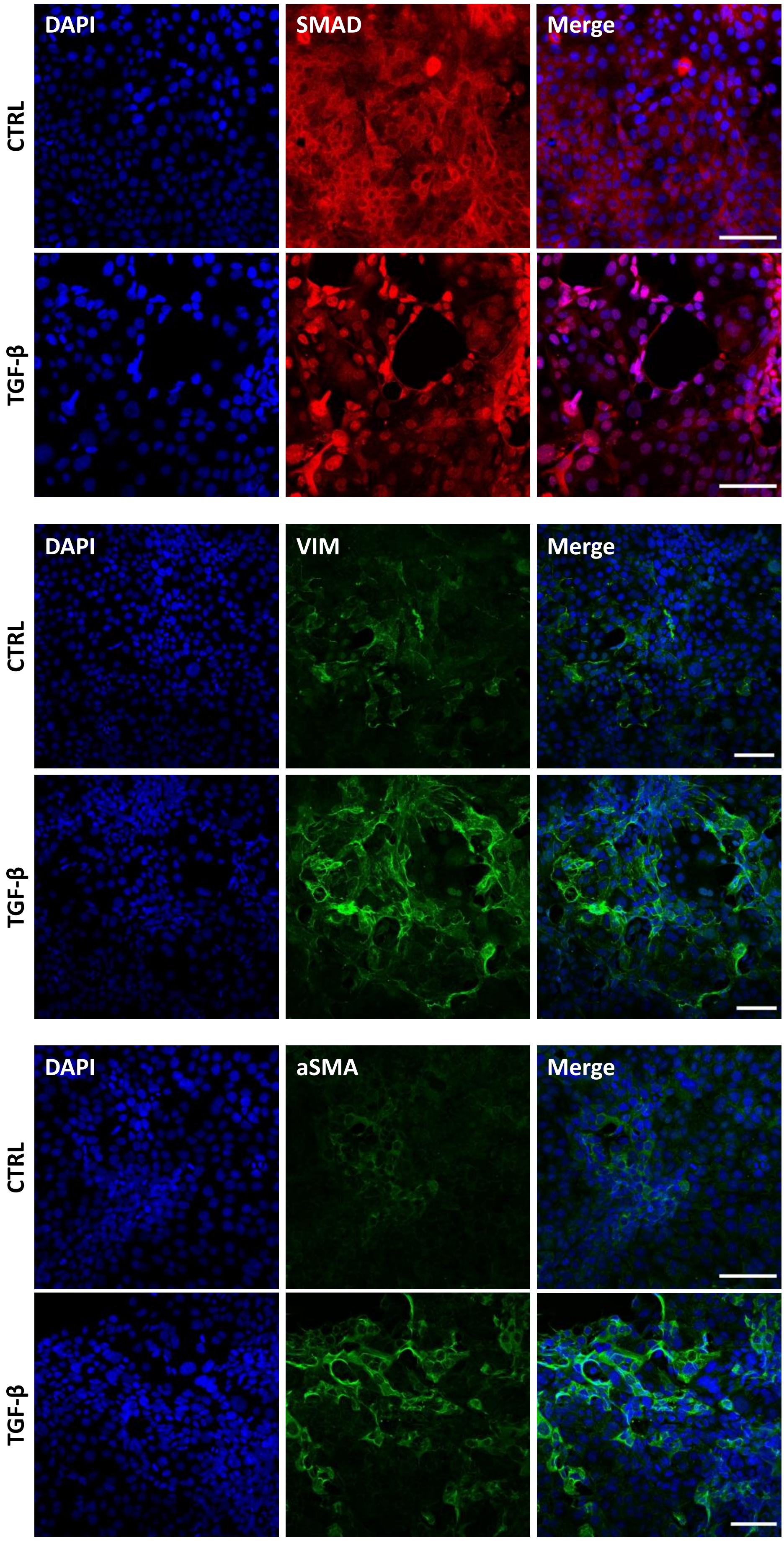
DAPI
SMAD
Merge
CTRL
TGF-β
DAPI
VIM
Merge
CTRL
TGF-β
DAPI
aSMA
Merge
CTRL
TGF-β

**Figure S6** – Representative confocal images of the expression of the indicated markers in PNT2 prostate cells treated or not with TGF-β (10 µg/mL) for 48 hours. CTRL refers to the control untreated cells. Scale bars: 100 µm.

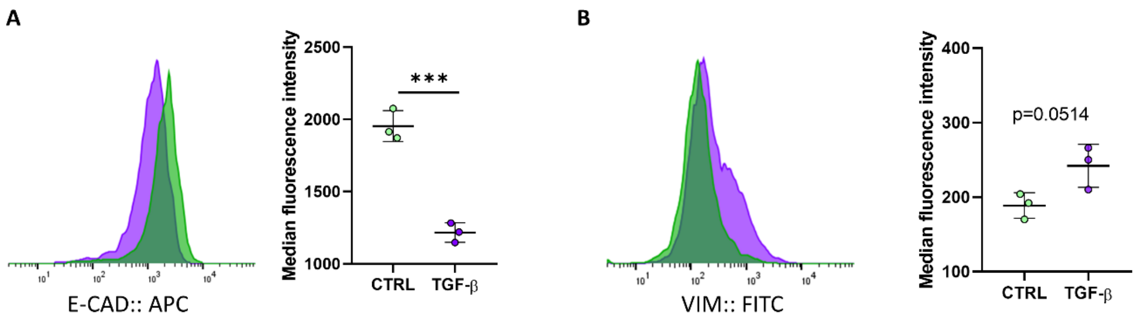


**Figure S7** – Flow cytometry quantification of (A) E-cadherin and (B) vimentin expression in PNT2 cells treated or not with TGF-β (10 µg/mL) for 48 hours. Statistical analysis performed by unpaired t-test.

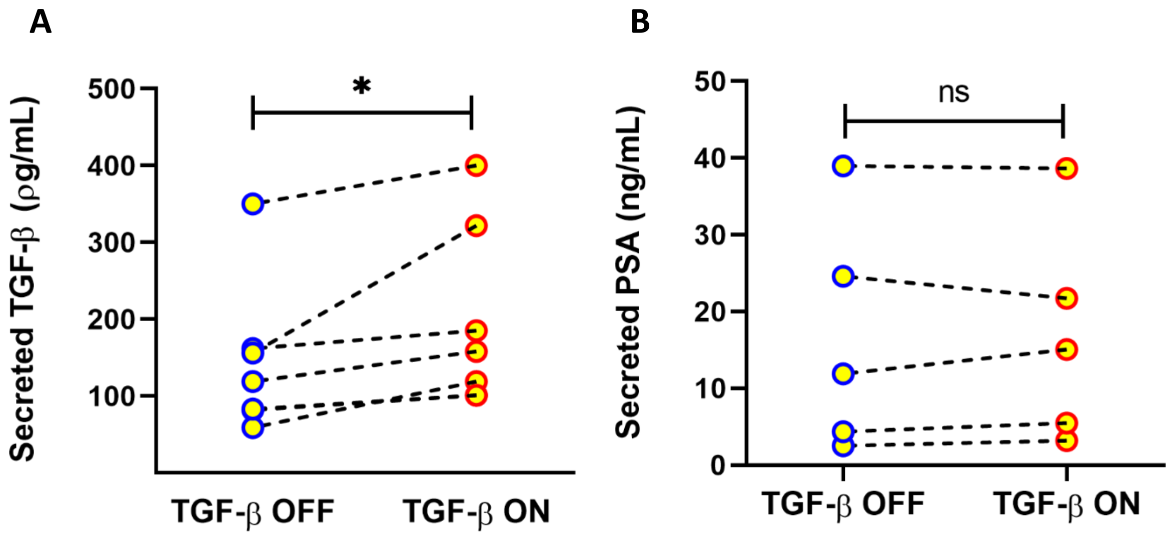


**Figure 8** – (A) Quantification of TGF-β secretion in PCTs isolated from 7 different patient tissues, as measured by ELISA. (B) Quantification of PSA secretion in PCTs isolated from patient tissues as measured by ELISA at day 5-8 of culture. Statistical analysis performed by paired t-test, n = 5.

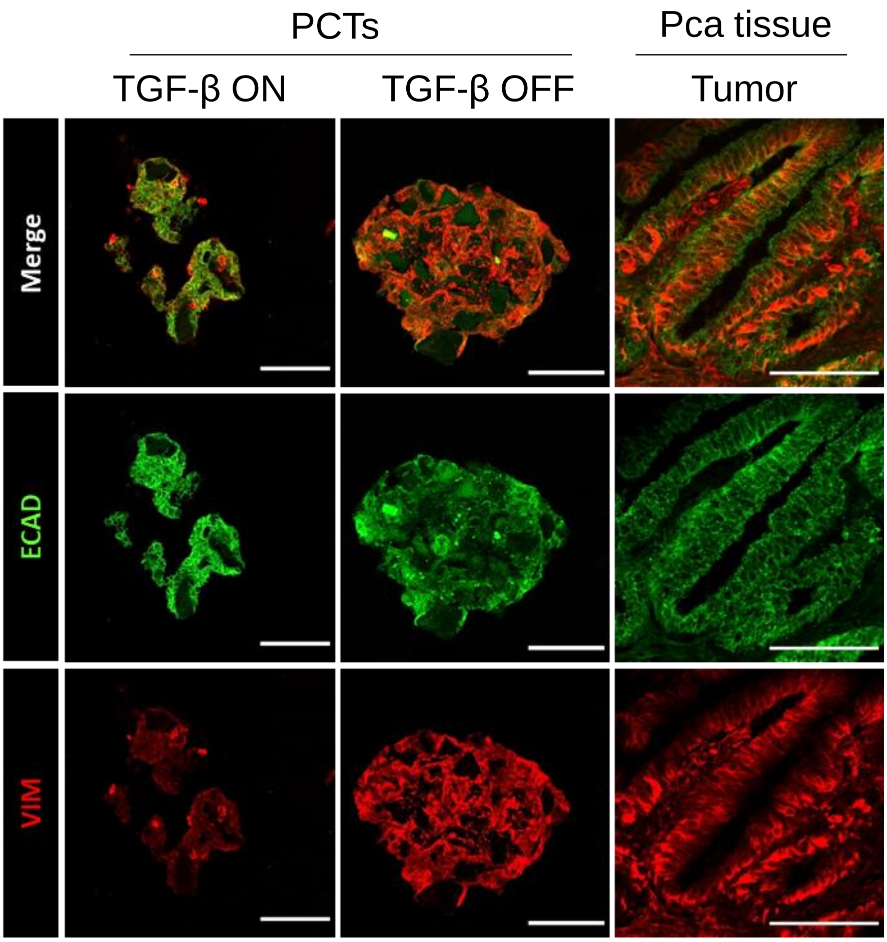


**Figure S9**. Representative confocal images of the expression of the indicated markers in PCTs. The PCTs were fixed with PFA 4 % and then embedded in OCT for slicing, prior to immunofluorescence staining. Scale bars: 50 µm.

**Supplementary Table 2.** RNA Sequencing raw data and significant regulated genes on PCTs TGF-β ON *vs* PCTs TGF-β ON.

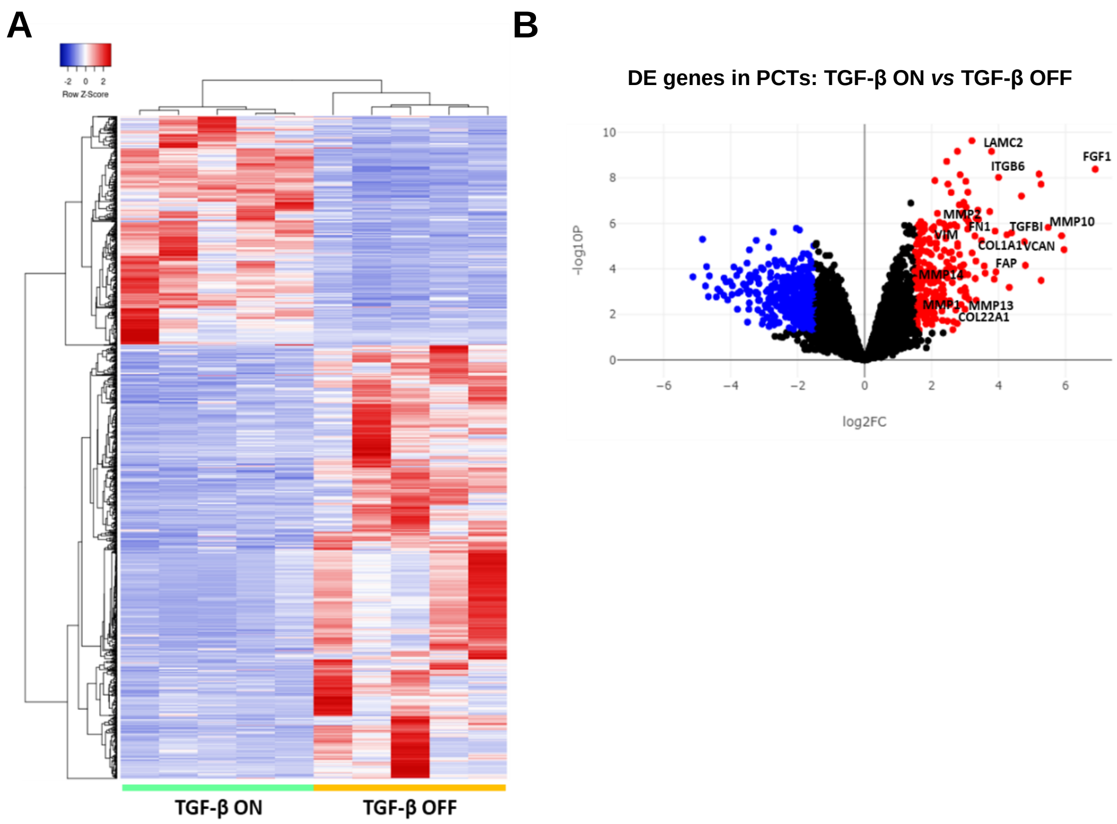


**Figure S10.** (A) Heatmap cluster and (B) Volcano plot representation of up and downregulated genes in TGF-β OFF and TGF-β ON PCTs (1.5-fold up or down from pooled normal, P adj < 0.05). n = 5.

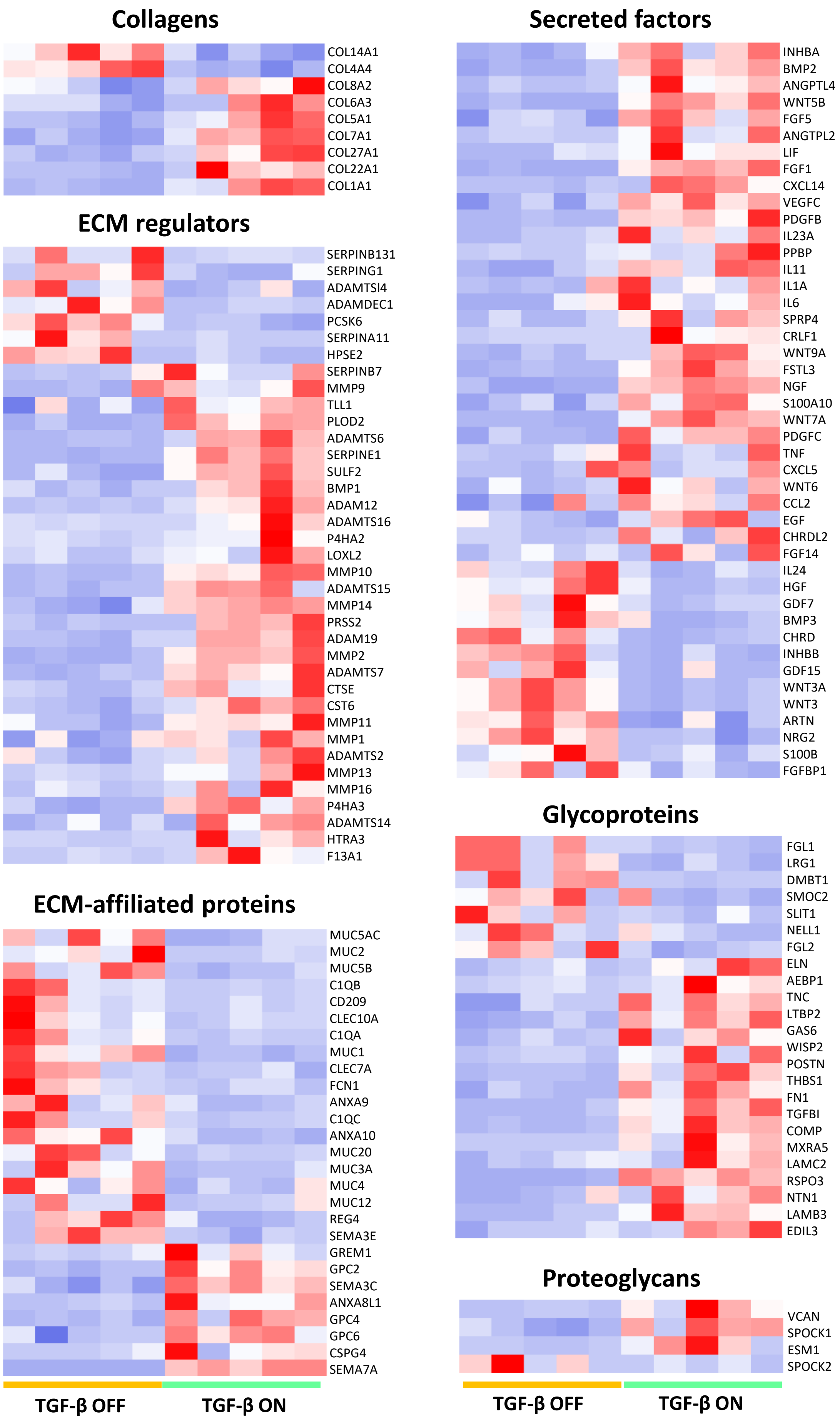
Collagens
COL14A1
COL4A4
COL8A2
COL6A3
COL5A1
COL7A1
COL27A1
COL22A1
COL1A1
ECM regulators
SERPINB131
SERPING1
ADAMTSl4
ADAMDEC1
PCSK6
SERPINA11
HPSE2
SERPINB7
MMP9
TLL1
PLOD2
ADAMTS6
SERPINE1
SULF2
BMP1
ADAM12
ADAMTS16
P4HA2
LOXL2
MMP10
ADAMTS15
MMP14
PRSS2
ADAM19
MMP2
ADAMTS7
CTSE
CST6
MMP11
MMP1
ADAMTS2
MMP13
MMP16
P4HA3
ADAMTS14
HTRA3
F13A1
ECM-affiliated proteins
MUC5AC
MUC2
MUC5B
C1QB
CD209
CLEC10A
C1QA
MUC1
CLEC7A
FCN1
ANXA9
C1QC
ANXA10
MUC20
MUC3A
MUC4
MUC12
REG4
SEMA3E
GREM1
GPC2
SEMA3C
ANXA8L1
GPC4
GPC6
CSPG4
SEMA7A
TGF-β OFF
TGF-β ON
Secreted factors
INHBA
BMP2
ANGPTL4
WNT5B
FGF5
ANGTPL2
LIF
FGF1
CXCL14
VEGFC
PDGFB
IL23A
PPBP
IL11
IL1A
IL6
SPRP4
CRLF1
WNT9A
FSTL3
NGF
S100A10
WNT7A
PDGFC
TNF
CXCL5
WNT6
CCL2
EGF
CHRDL2
FGF14
IL24
HGF
GDF7
BMP3
CHRD
INHBB
GDF15
WNT3A
WNT3
ARTN
NRG2
S100B
FGFBP1
Glycoproteins
FGL1
LRG1
DMBT1
SMOC2
SLIT1
NELL1
FGL2
ELN
AEBP1
TNC
LTBP2
GAS6
WISP2
POSTN
THBS1
FN1
TGFBI
COMP
MXRA5
LAMC2
RSPO3
NTN1
LAMB3
EDIL3
Proteoglycans
VCAN
SPOCK1
ESM1
SPOCK2
TGF-β OFF
TGF-β ON

**Figure S11**. Heatmap cluster representation of the 144 differentially expressed genes coding for matrisome proteins found significantly regulated in PCTs treated with TGF-β compared to non treated ones (1.5-fold, $P_{adj}$ < 0.05). n = 5.

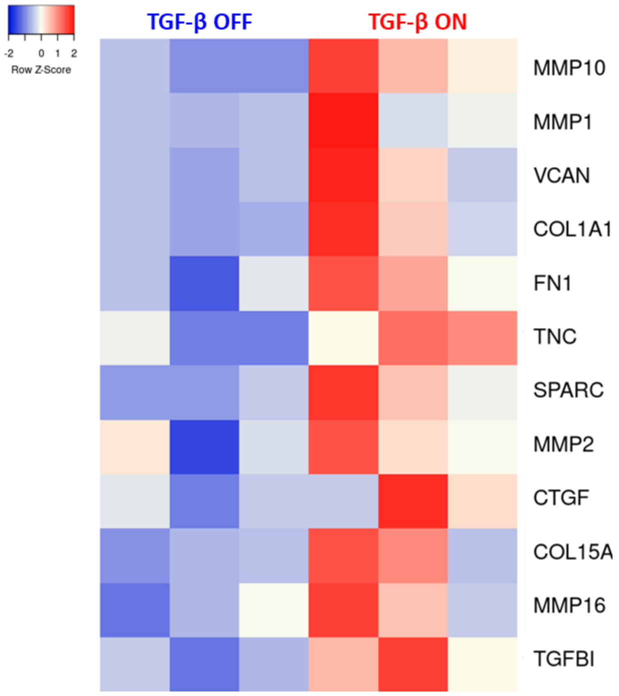


**Figure S12**. Heatmap cluster representation of the ECM genes found differentially expressed in PCTs PCTs treated with TGF-β compared to non treated ones as obtained by RT2 profiler PCR array ($P_{adj}$ < 0.05), n = 3.

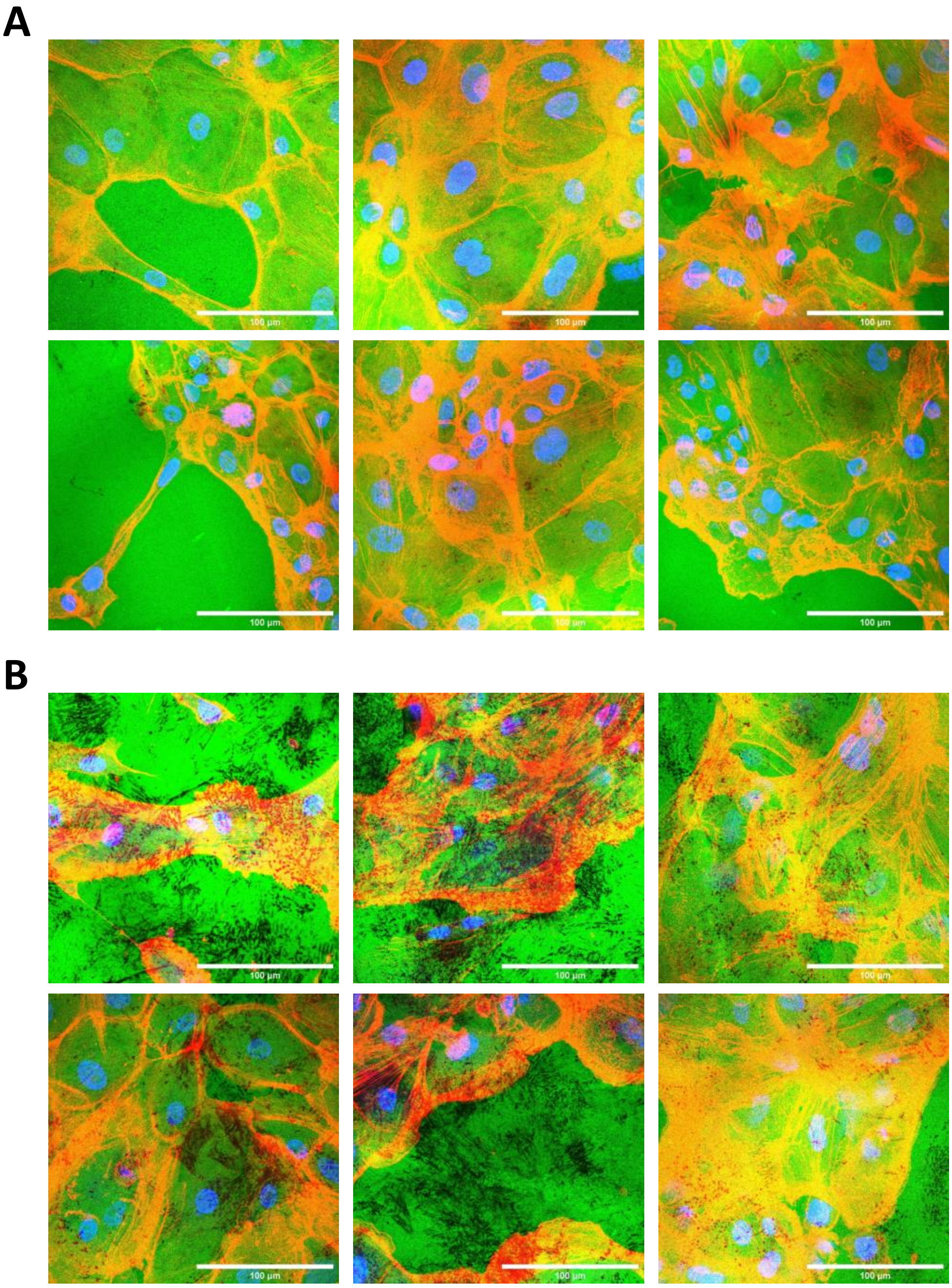


**Figure S13**. Representative confocal images of the gelatin degradation assay for cells with TGF-β OFF **(A)** and TGF-β ON **(B)**. Gelatin is green, F-actin is red and cell nuclei is blue DAPI. Scale bars: 100 µm.

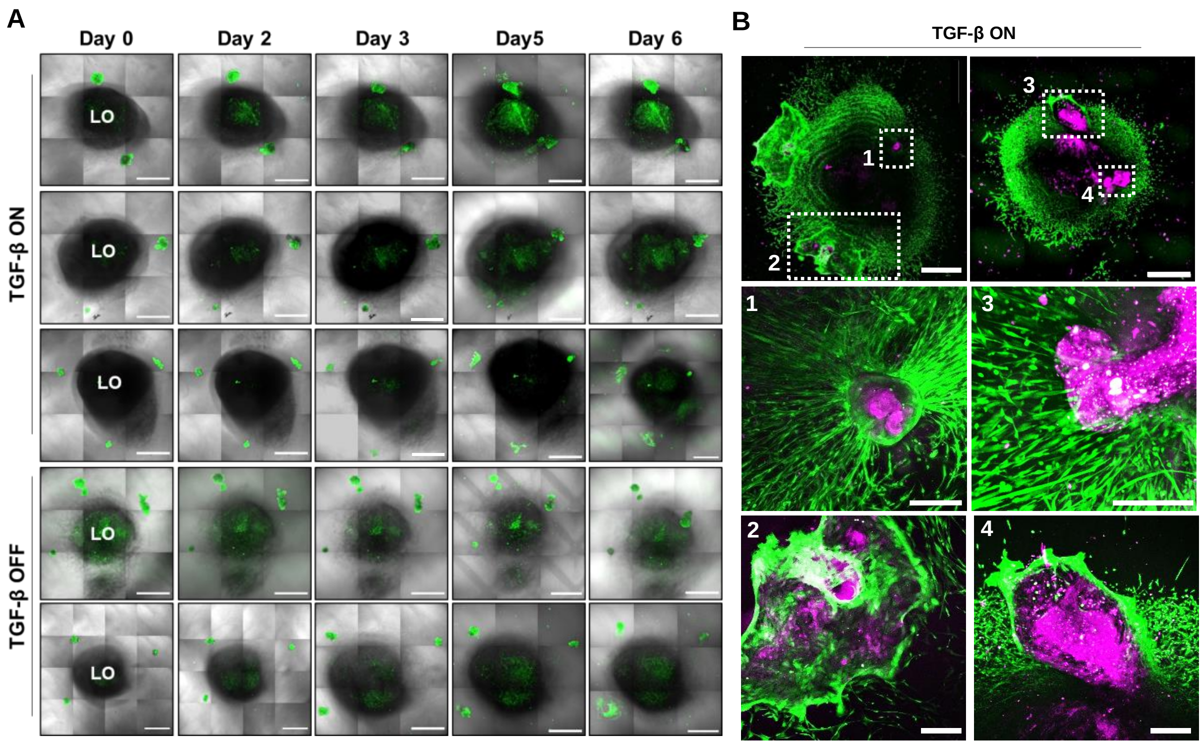


**Figure S14**. (A) Representative confocal live images of the confrontation assay performed using hiPSCs-derived lung organoids (LO) and PCTS (Cell tracker stained - green). Scale bars: 1000 µm Magnified brightfield images of PCTs. Scale bars: 1000 µm. **(B):** Representative live images from the confrontation assay, at day 7, for the PCTs TGF-β ON stained with CellTracker (pink) invading the LOs, with the whole culture stained for Calcein AM in green. Scale bars: 500 µm for the whole mounts and 200 µm for the magnified images 1-4.

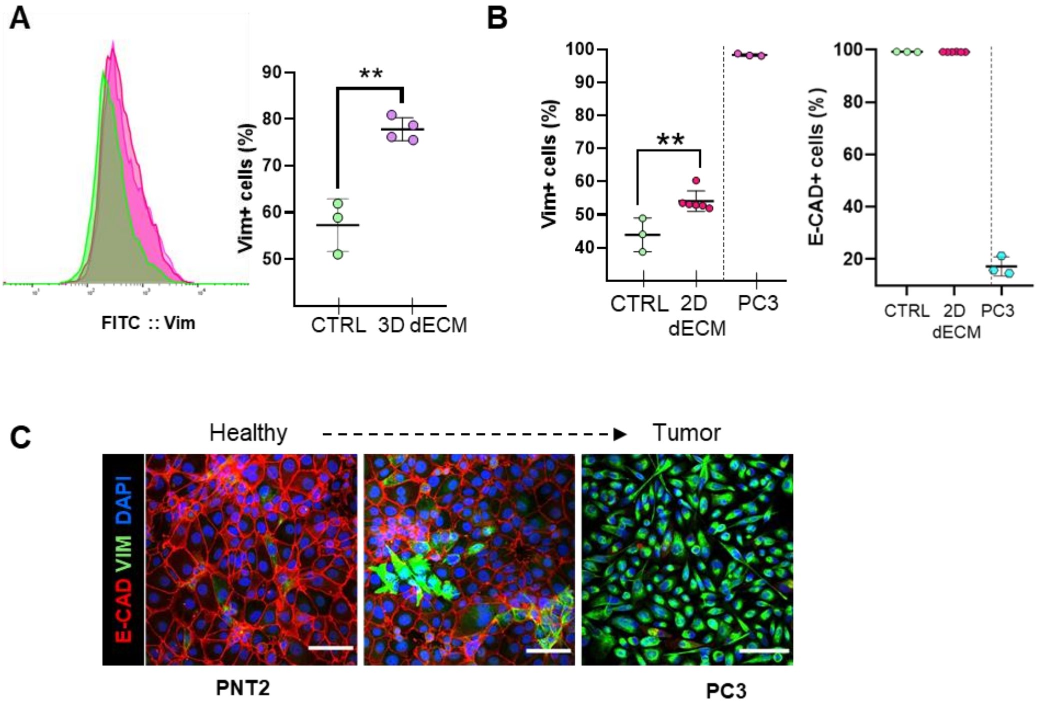


**Figure S15**. **(A)** Flow cytometry histogram representation and quantification of vimentin positive PNT2 cells, after a week of incubation with 3D dECMs (N = 2, n = 2, pink) and control cells grown in plastic (n = 3, green). Statistical analysis performed by t-test. **(B)** Flow cytometry quantification of vimentin and E-cadherin positive PNT2 cells, after 2 days of incubation in 2D dECM coated surface. PC3 are used as positive control for tumour cells expressing high levels of vimentin and low levels of E-cadherin. Statistical analysis was done by one-way ANOVA **(C)** Representative confocal images of PNT2 and PC3 cells, stained with E-cadherin (red) and vimentin (green). Scale bars: 100 µm.
